# Consensus mutagenesis and ancestral reconstruction provide insight into the substrate specificity and evolution of the front-end Δ6-desaturase family

**DOI:** 10.1101/2020.02.07.938332

**Authors:** Dongdi Li, Adam M. Damry, James R. Petrie, Thomas Vanhercke, Surinder P. Singh, Colin J. Jackson

**Affiliations:** Research School of Chemistry, The Australian National University, Canberra, ACT 2601, Australia; CSIRO Agriculture Flagship, Black Mountain Laboratories, Canberra, ACT 2601, Australia

**Author notes:** These authors contributed equally to this work (Dongdi Li, Adam M. Damry). To whom correspondence should be addressed: Colin J. Jackson, Research School of Chemistry, The Australian National University, ACT 0200, Australia.

## Abstract

Marine algae are a major source of omega (ω)-3 long-chain polyunsaturated fatty acids (ω3-LCPUFAs), which are conditionally essential nutrients in humans and a target for industrial production. The biosynthesis of these molecules in marine algae begins with the desaturation of fatty acids by Δ6-desaturases and enzymes from different species display a range of specificities towards ω3 and ω6 LCPUFAs. In the absence of a molecular structure, the structural basis for the variable substrate specificity of Δ6-desaturases is poorly understood. Here we have conducted a consensus mutagenesis and ancestral protein reconstruction-based analysis of the Δ6-desaturase family, focusing on the ω3-specific Δ6-desaturase from *Micromonas pusilla* (MpΔ6des) and the bispecific (ω3/ω6) Δ6-desaturase from *Ostreococcus tauri* (OtΔ6des). Our characterization of consensus amino acid substitutions in MpΔ6des revealed that residues in diverse regions of the protein, such as the N-terminal cytochrome b5 domain, can make important contributions to determining substrate specificity. Ancestral protein reconstruction also suggests that some extant Δ6-desaturases, such as OtΔ6des, could have adapted to different environmental conditions by losing specificity for ω3-LCPUFAs. This dataset provides a map of regions within Δ6-desaturases that contribute to substrate specificity and could facilitate future attempts to engineer these proteins for use in biotechnology.

## INTRODUCTION

Omega (ω)-3 long-chain polyunsaturated fatty acids (ω3-LCPUFAs) such as docohexaenoic acid (DHA, C22:6^Δ4,7,10,13,16,19^) and eicosapentaenoic acid (EPA, C20:5^Δ5,8,11,14,17^) are conditionally essential fatty acids in humans that are vital to healthy metabolism.^*1*^ Deficiencies in these dietary fatty acids results in a reduction of cell membrane fluidity that can adversely affect neurotransmission and contribute to cardiovascular disease.^*2, 3*^ However, most animals have a limited capacity to synthesize ω3-LCPUFAs as they lack key desaturase enzymes that catalyze the production of ω3-LCPUFAs from precursors such as linoleic acid (LA, C18:2^Δ9,12^) and α- linolenic acid (ALA, C18:3^Δ9,12,15^) by introducing double bonds into fatty acid acyl chains. In nature, marine algae are some of the primary producers of ω3-LCPUFAs *via* the Δ6-desaturase-dependent pathway, which then accumulate in fish oils that serve as a major dietary source of ω3-LCPUFAs for humans. However, due to the scarcity of marine resources, relying wholly on fish oils for dietary ω3-LCPUFA requirements is projected to be unsustainable. More efficient methods of ω3-LCPUFA production are thus of interest, the development of which requires a better understanding of the enzymes that catalyze the rate limiting commitment step in their synthesis: Δ6-desaturases.^*4-7*^

Δ6-desaturases belong to the family of membrane-bound front-end desaturases, which catalyze double bond formation at specific positions near the fatty acid carboxyl end. Δ6-desaturases specifically introduce a double bond between the acyl chain C6 and C7, and preferentially target substrates such as LA and ALA that already possess several double bonds including at the Δ9 position.^*4, 8*^ Although crucial to LCPUFA-production pathways, these enzymes have proven to be extremely difficult to characterize in depth. Very few high-resolution structures of membrane-bound fatty acid desaturases have been solved, none of which are of Δ6-desaturases, and these enzymes cannot be easily purified in an active form. Sequence-based modelling and topology studies have nonetheless provided cursory information about the organization of these enzymes, generally predicting a four-transmembrane helix (TMH) topology that resembles that of the few crystallized Δ9-desaturases.^*9-11*^ These TMHs form one portion of the enzymes’ desaturase domain, which also contains the enzyme active site and three conserved histidine-rich motifs (His boxes) that are thought to coordinate a di-iron center.^*12-15*^ Δ6-desaturases are also known to require a fused b5 cytochrome domain at their N-terminus for their activity, which functions as an electron donor during catalysis.^*16-18*^ However, although several other desaturases are capable of accepting electrons from cytoplasmic cytochromes, mutating key residues in the b5 domain of Δ6-desaturases renders the enzymes non-functional, hinting that this domain may serve additional roles in Δ6-desaturases.^*19, 20*^

The LCPUFA Δ6-desaturase-dependent biosynthesis pathway was recently reconstructed in model and crop plant systems using the *Micromonas pusilla* Δ6-desaturase (MpΔ6des) (Supplementary Fig. 1).^*4, 5, 21, 22*^ MpΔ6des was chosen to catalyze this reaction, because it is among the most active ω3 specific Δ6-desaturase to be identified from algae or plant.^*4, 6*^ Other Δ6-desaturases, such as that from *Ostreococcus tauri* (OtΔ6des), do not display a preference for ω3-LCPUFAs and are therefore less desirable for biotechnology applications.^*23, 24*^ Although MpΔ6des is an efficient and ω3-specific Δ6-desaturase, further optimization of this enzyme, particularly in terms of its specificity for ω3-substrates such as α-linolenic acid, would result in enhanced recombinant LCPUFA production in plants. Mutagenesis studies have revealed that the second putative TMH, the regions adjacent to the conserved histidine motifs and the C-terminus affect the regioselectivity and the preferred substrate chain length of membrane desaturases.^*10, 11, 25, 26*^ However, although Δ6-desaturases have been identified that naturally prefer either ω3 or ω6-fatty acids,^*23, 27-31*^ detailed mutagenesis studies to modify ω3/ω6-fatty acid preference have not been performed, to the best of our knowledge. Similarly, the evolution of ω3/ω6-fatty acid specificity, including divergence between MpΔ6des and OtΔ6des, has not been investigated in detail.

**Figure 1.**
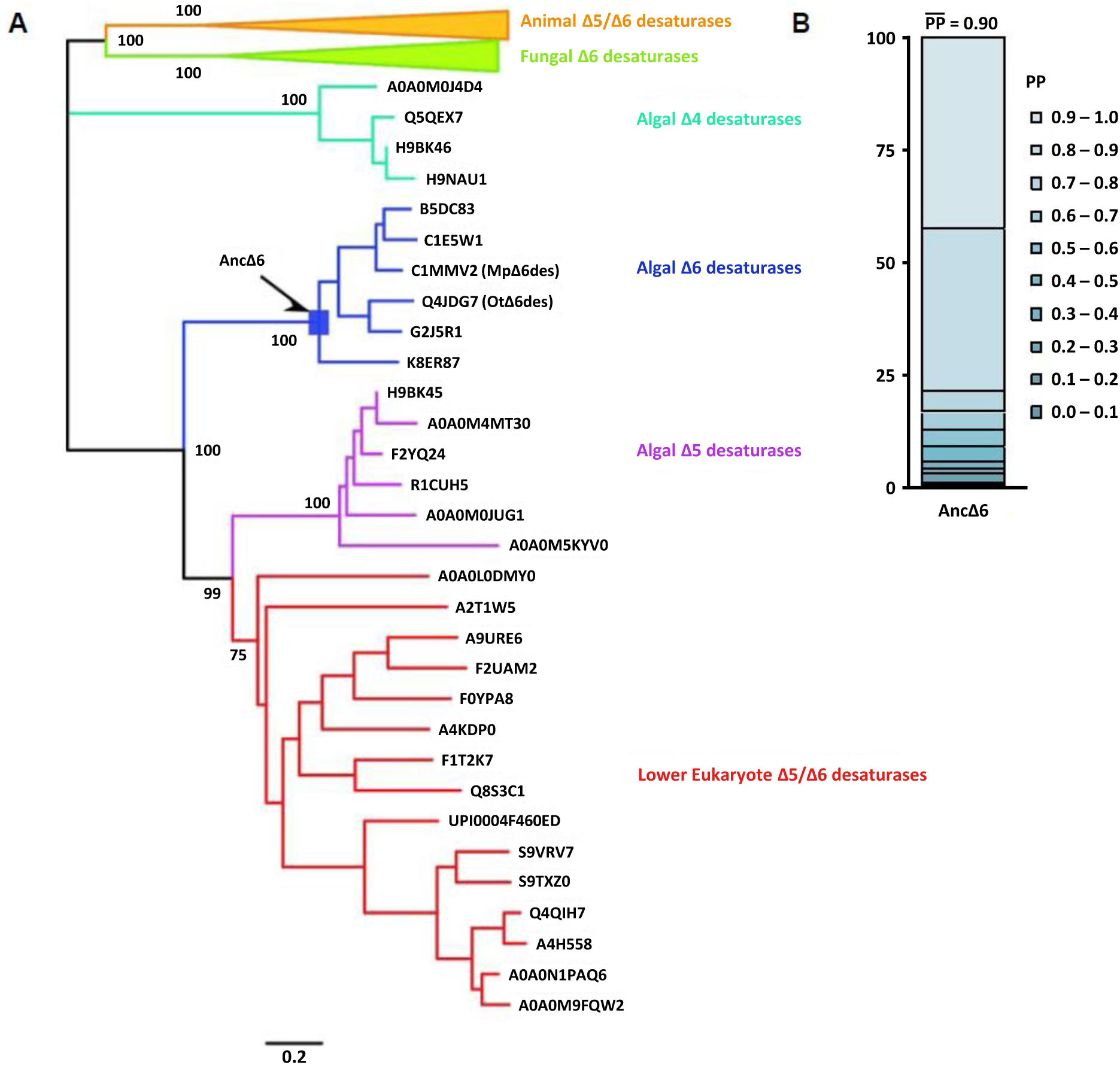
Phylogenetic analysis of desaturases. **A.** An unrooted phylogeny tree was constructed using MEGA v.6. The analysis demonstrated the clear phylogenetic separation of algal, fungal, and animal desaturases. Included desaturase sequences are included in Supplementary Table 1. The bootstrap values (100 bootstrap repetitions) of major nodes are denoted. The identity of the protein at each node is presented using its UniProt ID with MpΔ6des and OtΔ6des denoted for clarity. **B**. Posterior probability distribution and mean posterior probability of the inferred Δ6-desaturase ancestor AncΔ6.

Here, we have conducted an intensive mutagenesis study aimed at understanding Δ6-desaturase specificity through the introduction of consensus mutations from the larger algal desaturase family and ancestral protein reconstruction. This study identifies several regions of the protein that affect substrate preference, including a more significant role for the cytochrome b5 domain than previously thought. These results combined with modelling and topology predictions also suggest mechanisms by which desaturases gate substrate specificity. Overall, this work takes important steps to elucidate the complexities of desaturase substrate specificity, laying a foundation for the future engineering of these critical enzymes along the ω3-LCPUFA production pathway.

## METHODS

### Yeast vectors and strains

An isogenic strain of *Saccharomyces cerevisiae* S288C (*his3D200, ura3-52, leu2D1, lys2D202 and trpD63*)^*32*^ was used throughout this project for the expression of wild-type and variants of MpΔ6des. The pYES2-ura yeast expression vector containing the wild-type MpΔ6des gene was used for the expression and mutagenesis of the desaturases. *Escherichia coli* TOP10 cells were used for plasmid preparation and cloning procedures.

### Phylogenetic analysis and structural modelling of algae Δ6-desaturases

MpΔ6des (UniProt ID: C1MMV2) was used as the query sequence and BLASTP was used as the search engine to retrieve 100 homologues of front-end desaturases in the non-redundant protein sequence database.^*33, 34*^ After the sequence set was processed using the CD-HIT suite^*35*^ to remove sequences with more than 90% sequence identity, duplicate sequences and highly similar sequences were removed. The number of sequences was reduced to 57, which were aligned with the MUSCLE algorithm^*36*^ in the MEGA6 program.^*37*^

A phylogenetic tree was constructed using the maximum-likelihood method implemented by the MEGA6 program.^*37*^ The phylogenetic tree was constructed using the maximum likelihood method with LG rate matrix, and bootstrapped with 100 replicates. The ancestral protein AncΔ6 was constructed from the phylogenetic tree using the empirical Bayesian method that is implemented in the PAML4 software package.^*38*^ Because of the low sequence conservation of the N-terminus, the fragment 1-54 of wild-type MpΔ6des replaced the low-confidence inferred N-terminus of the ancestor.

The transmembrane topology prediction was carried out using the CCTOP webserver^*39*^ and the visual representation generated using the PROTTER webserver.^*40*^ Amino acid sequences for MpΔ6des, OtΔ6des, and AncΔ6 were then submitted to the Phyre2 webserver for homology protein structure prediction.^*41*^ Desaturase domains for all three were modelled using the two available acyl-CoA desaturase structures (PDB ID: 4YMK^*42*^ and 4ZYO^*43*^) as templates and cytochrome b5 domains were modelled using several free cytochrome b5 structures as templates.

### Construction of MpΔ6des Variants

The previously constructed pYES2–ura plasmid with the MpΔ6des gene cloned between *Kpn*I and *Sac*I sites was used as the template to generate the desired single mutants. Primer pairs were designed according to the QuikChange site-directed mutagenesis protocol.^*44*^ A T7 primer was used with the anti-sense primer and Gal1 terminator primer was used with the sense primer of each mutagenesis primer pair to generate DNA fragments with the desired mutation in the overlapping region. The DNA fragments and the *Kpn*I/*Sac*I-digested pYES2 vector were assembled using an in-house Gibson assembly kit.^*45*^ The resulting coding regions were confirmed by sequencing. The 23 algal Δ6 consensus mutants were constructed and cloned by GenScript.

### Functional expression in yeast

Yeast expression plasmids were transformed into *S. cerevisiae* S288C using the Yeast-1 transformation kit (Sigma Aldrich, Australia), according to the provided protocol. Transformants were screened on yeast minimal growth agar supplemented with 2% agar, 2% glucose and an amino acid mixture lacking uracil (SC^-^ -Ura Glu). Successful transformants were confirmed by colony PCR.

Starter cultures of S288C transformants were set up in 5 ml of SC^-^ -Ura medium supplemented with 2% glucose. After incubation overnight at 28 °C with shaking, yeast cells were washed with sterile deionized water. The washed yeast cells were resuspended in 5 ml of SC^-^-Ura medium supplemented with 1% Tergitol, 2% galactose, 0.25 mM of α-linolenic acid (C18:3n-3^Δ9, 12, 15^) and 0.25 mM of linoleic acid (C18:2n-6^Δ9, 12^) at OD600 of approximately 0.3. After overnight incubation at 28 °C with shaking, the yeast cells were washed successively with 5 ml of 1% Tergitol, 0.5% Tergitol, and deionized water to remove non-incorporated free fatty acids. The yeast pellets were freeze-dried for lipid analysis.

### HA tagging of wild-type MpΔ6des and its variants

Primers were designed to add an influenza haemagluttinin (HA) tag onto the N-terminus of MpΔ6des (Table 2) via blunt-end ligation. The N-terminal HA tagged variants were generated by assembling the N-terminus of the tagged WT MpΔ6des with the variant genes and the *Kpn*I/*Sac*I-digested pYES2 vector using an in-house Gibson assembly kit. The tagged coding regions were confirmed by sequencing.

### Lipid analysis

Fatty acid methyl esters were prepared from the freeze-dried yeast pellets by transesterification with 1 ml methylation solution (MeOH : dichloromethane : HCl = 10 : 1 : 1) for 2 hours at 80 °C. After adding 1 mL of deionized water into each sample, the methylated fatty acids were extracted with 0.3 mL of extraction solution (hexane : dichloromethane = 4 : 1) and reconstituted in hexane before GC analysis.^*4*^ The desaturation efficiency was calculated based on the integrated peak area as described previously^*4*^ and normalized using the wild-type desaturase result in the same assay set.

### Western blot analysis

The S288C transformant cultures were prepared as for the feeding assay, except that Tergitol and fatty acid substrates were not added into the induced culture. The yeast cell pellets were washed with 1 mL 0.1 M Tris-HCl (pH 7.2). The yeast lysate was prepared using 0.5 mm zirconium oxide beads and a Bullet blender (Next Advance, Cambridge, MA) in 200 μL resuspension buffer (50 mM Tris-HCl pH 7.9, 20 mM glycerol, 1 mM DTT). The total protein concentration was determined using the Bradford Protein Assay Kit (Thermo Scientific, Waltham, MA) with albumin as a standard for each western blot.

A sample of protein lysate equivalent to 40 µg of total protein was run on a NuPAGE 10% Bis-Tris Protein Gel (NOVEX, Thermo Fisher Scientific) and transferred to a piece of nitrocellulose membrane using a semi-dry transfer apparatus (Hoefer ® Semiphor^TM^, Pharmacia). The membrane was blocked for 1 hour with blocking buffer (5 % skim milk suspension in 20 mM Tris-HCl, 8% NaCl, 0.1 % Tween 20, pH 7.6) and probed with anti-HA antibody at 1:5000 dilution (Sigma) in blocking buffer. The membrane was then washed 4 times with 20 mL TBST (20 mM Tris-HCl, 8% NaCl, 0.1 % Tween 20, pH 7.6) for 10 minutes and probed with anti-mouse horseradish peroxidase conjugated antibody at 1:10000 dilution in blocking buffer. The membrane was washed 4 times again as before and 1:1 mixture of peroxide solution and the luminol/enhancer solution from the SuperSignal®chemiluminescent substrate kit (Pierce) was added. The blot was visualized with ChemiDoc (Bio-Rad).

## RESULTS

### Phylogenetic analysis of MpΔ6des homologs

In the absence of any molecular structure to guide mutagenesis, we first sought to obtain deeper insight into these proteins through phylogenetic analysis. Previous phylogenetic studies of membrane-bound desaturases revealed the evolutionary separation of acyl-CoA desaturases and acyl-lipid desaturases.^*4*^ For the purpose of consensus mutation design and ancestral protein reconstruction (APR), we conducted a focused phylogenetic analysis of MpΔ6des homologs. The protein basic local alignment search tool (BLAST) was used to search the non-redundant protein database with the MpΔ6des protein sequence.^*34*^ A cutoff of 26% sequence identity to MpΔ6des was applied to obtain a dataset of closely related sequences containing Δ4-, Δ5- and Δ6-desaturases from animals, simple unicellular organisms and marine green algae (Supplementary Table 1). No close homologs were found in modern plants. The phylogenetic analysis of the above members using a maximum likelihood algorithm showed the clear separation of animal desaturases from the Δ5- and Δ6-desaturases of other organisms (Fig. 1). In each group, there are separate Δ5-desaturase and Δ6-desaturase clades. Algal Δ6-desaturases form one clade, whereas the Δ5 clade contains Δ5-desaturases from algae and a number of other related unicellular organisms.^*46*^ Δ4-desaturases are only present in algae and form a distinct clade from the other algal desaturases. Considering that Δ4-desaturation is the last step of ω3-LCPUFA biosynthesis that converts DPA to DHA (Supplementary Fig. 1), the clustering of algal Δ4-desaturases may indicate a later gene duplication event as part of cold-adaptation, since the increased desaturation of the C22 carbon chain could result in higher membrane fluidity.^*47-50*^ Despite the evolution of land plants from an algal common ancestor, the rare higher plant Δ6-desaturases are thought to have originated through retrotranscription, rather than by direct inheritance.^*51*^ Hence, the absence of a modern plant acyl-CoA Δ6-desaturase homologue is not surprising.

### Topology prediction of MpΔ6des

Although the algal Δ6-desaturases have been functionally characterized, we understand little about how their function relates to their structure.^*4, 28, 30*^ Only two classes of membrane-bound desaturase topologies have been experimentally verified. Most eukaryotic desaturases adopt a four TMH topology, including the *Mus musculus* and *Homo sapiens* Δ9 acyl-CoA desaturases (MmΔ9des and HsΔ9des, respectively), which are the only two membrane-bound desaturases for which high-resolution crystal structures have been solved (PDB ID: 4YMK and 4ZYO, respectively).^*42, 52*^ A second topology, first validated in the *Bacillus subtilis* Δ5 acyl-lipid desaturase (BsΔ5des), possesses six TMHs instead.^*53*^ Although these three proteins almost certainly evolved from a common ancestor and share conserved functional motifs that include their active site and the three His boxes, all three show substantial sequence variation to MpΔ6des, ranging from 16% to 20% identity (Supplementary Fig. 2). To investigate the structure of MpΔ6des and its interaction with the membrane, we performed a transmembrane helix prediction using the CCTOP topology prediction server,^*39*^ which predicts membrane protein topology via the consensus of ten different topology prediction models enhanced with available structural information from the TOPDB protein topology database.

**Figure 2.**
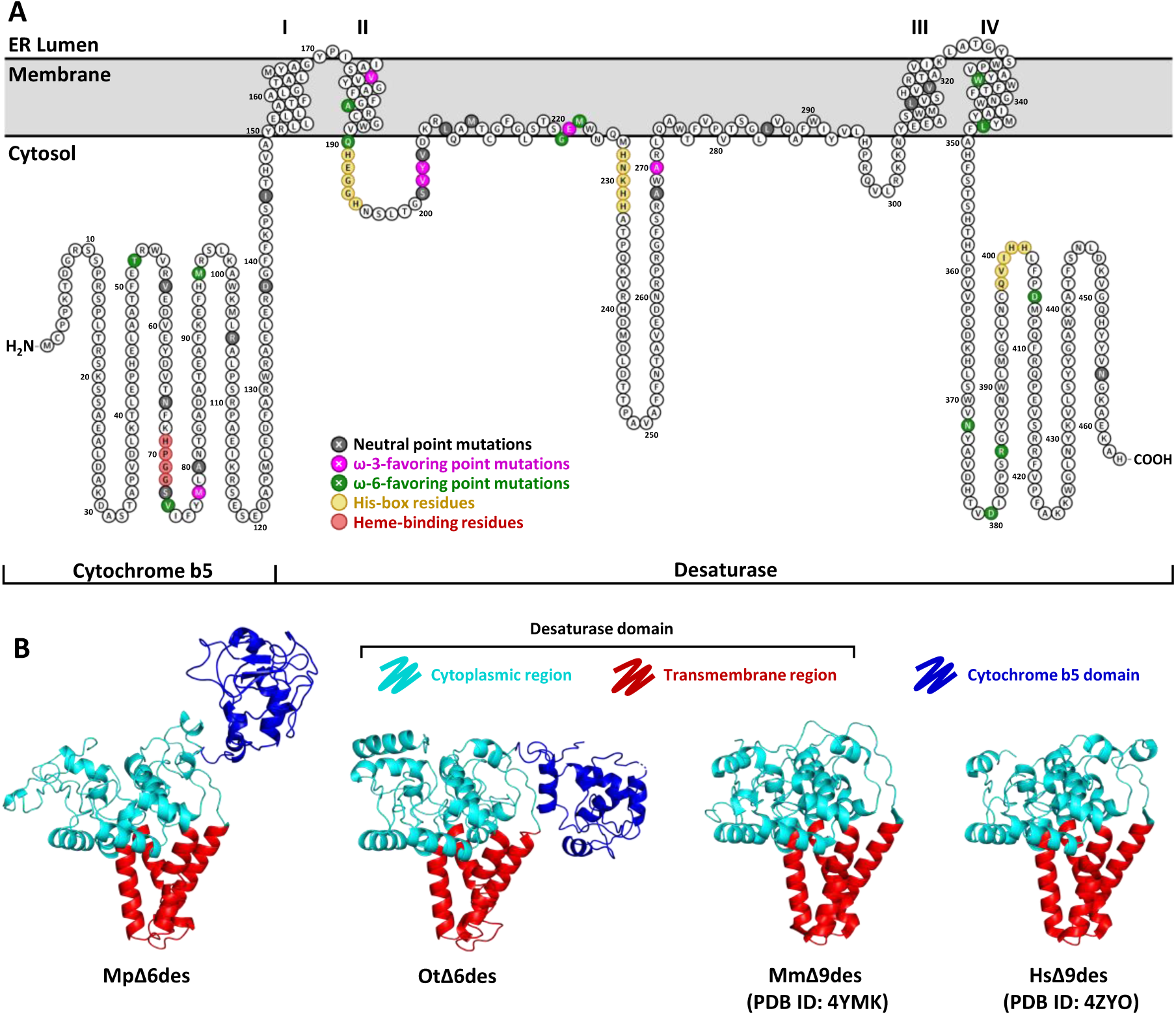
Predicted topology of MpΔ6des. **A.** The corrected 4 TMH topology for algal Δ6- desaturases is shown as a PROTTER topology schematic. The N- and C-termini as well as the His-box motifs indicated are known to be cytoplasmic. **B.** Phyre2 Models of MpΔ6des and OtΔ6des are shown side by side to the crystal structures of MmΔ9des and HsΔ9des. All four desaturases adopt a 4 TMH topology. However, MmΔ9des and HsΔ9des do not possess a cytochrome b5 domain, therefore despite the correct identification of the MpΔ6des and OtΔ6des b5 domains by the Phyre2 algorithm, their exact position relative to the linked desaturase domain in each protein is unknown.

The CCTOP consensus predicts three putative TMHs in MpΔ6des. However, unlike control predictions made for MmΔ9des and BsΔ5des, which are accurately predicted as possessing 4 TMHs and 6 TMHs respectively across the majority of the modeling methods and cross-references used by CCTOP, agreement between models for MpΔ6des is weaker, with the ten methods predicting between 3 and 7 TMHs (Supplementary Fig. 3). In addition, the proposed three TMH consensus cannot be accurate, as all three His boxes (positions 191-195, 228-232, and 398-402) and both termini are known to be cytoplasmic.^*10, 11, 46, 54*^ Across all ten models as well as in the consensus topology however, TMHs are predicted to occur in roughly the same positions as in MmΔ9des. It is therefore likely that the true topology of MpΔ6des possesses four TMHs, with the predicted TMH1 helix instead forming adjacent TMH1 and TMH2 helices similarly to MmΔ9des and HsΔ9des (Fig. 2a). We then used the Phyre2 webserver^*41*^ to generate homology models of MpΔ6des and OtΔ6des (Fig. 2b). Both resulting models utilize the existing Δ9 desaturase and cytochrome b5 crystal structures, further suggesting homology to these 4 TMH desaturases, and place the four MpΔ6des TMHs at positions 150-169, 173-189, 305-323, and 329-349.

**Figure 3.**
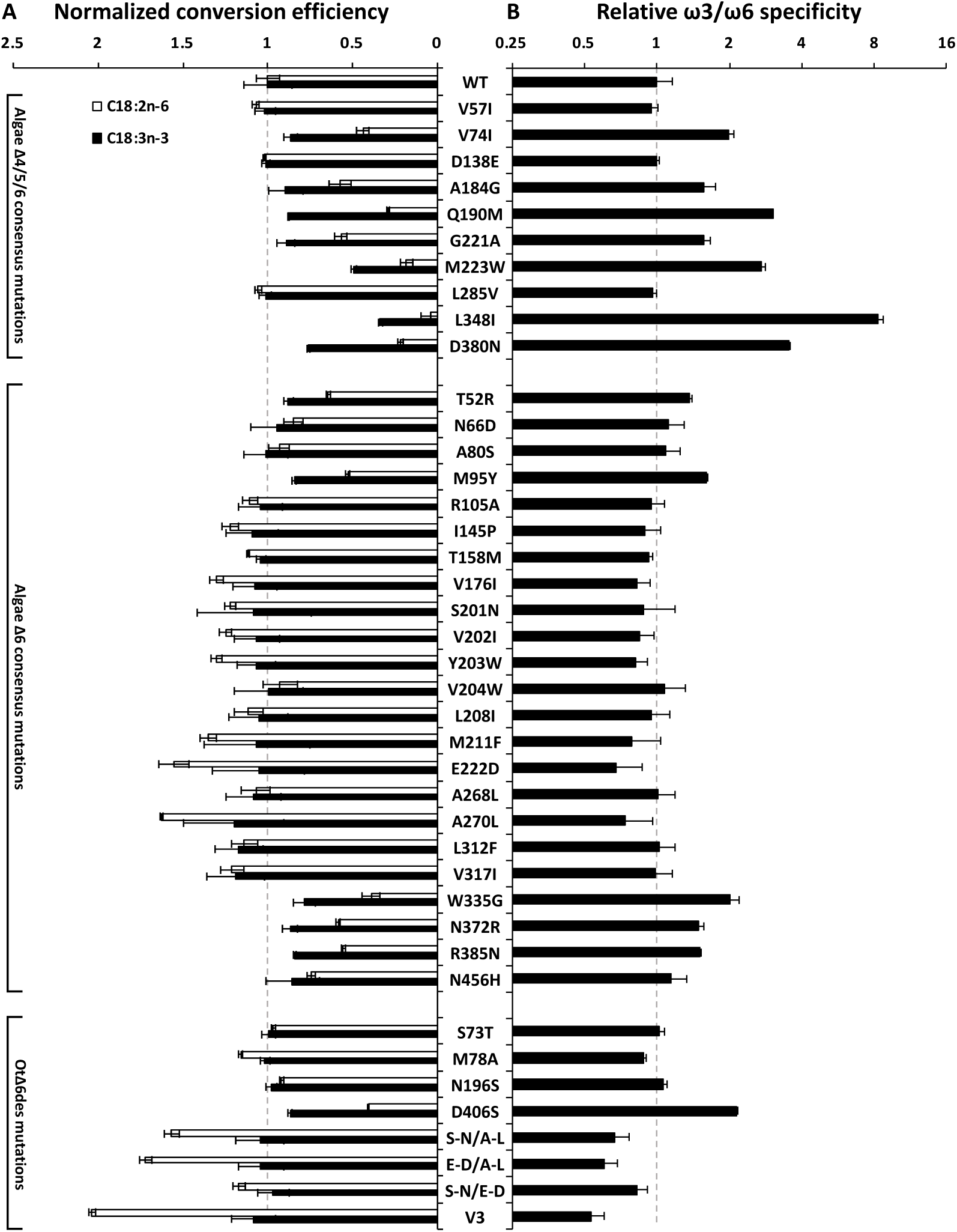
Activity and specificity of MpΔ6des consensus variants. **A.** Conversion efficiencies for both ω3 and ω6 substrates (ALA, black; and LA, white respectively) are shown for the MpΔ6des variants identified by consensus mutagenesis, normalized to wild-type conversion efficiency (n = 3 ± 1 S.D.). Results are grouped by the consensus set in which they were predicted. Combinatorial mutants, grouped with OtΔ6des, are named by the mutations included, with S-N representing the S201N mutation, E-D the E222D mutation, A-L the A270L mutation, with the exception of V3, which includes all three above mutations. **B.** Relative ω3/ω6 specificities for all variants are shown, normalized to the ω3/ω6 ratio of the wild type (n = 3 ± 1 S.D.). A ratio > 1 represents an increase in ω3 specificity, whereas a ratio < 1 represents an increase in ω6 specificity, relative to the wild type MpΔ6des.

### Identifying determinants of MpΔ6des specificity through consensus mutagenesis

Δ6- desaturases from different organisms are known to catalyze Δ6-desaturation of different acyl chains, with MpΔ6des for example exhibiting high specificity for ω3 substrates whereas OtΔ6des demonstrates comparable activity on both ω3 and ω6 fatty acids. Rationally inferring and engineering the determinants of desaturase specificity is however a difficult task given experimental limitations when working with desaturases. A lack of available high-resolution structural information prevents the use of computational design tools to predict specificity-linked residues in Δ6-desaturases, and the low-throughput nature of existing activity assays for these enzymes precludes the use of specificity-altering methodologies that require the screening of a large number of mutants such as directed evolution. We thus turned to consensus mutagenesis to locate positions responsible for MpΔ6des substrate specificity, as high probability mutations in the desaturase family are likely to be tolerated in the context of the MpΔ6des scaffold and are more likely to encode specificity information than random mutations.

For this purpose, we used three different amino acid consensus sets to identify suitable mutation sites. The first, based on an alignment of algal Δ4-, Δ5- and Δ6-desaturases (including the Δ5-desaturases from other unicellular organisms,) identified ten residues that are highly conserved throughout the alignment but differ in MpΔ6des (Supplementary Fig. 4). The second, based on an alignment of the six identified algal Δ6-desaturases (Supplementary Fig. 5) identified a further 23 residues that are highly conserved but not present in MpΔ6des. Finally, to investigate the difference in specificity between MpΔ6des and OtΔ6des, as OtΔ6des is the characterized algal Δ6-desaturase whose substrate specificity is most different to that of MpΔ6des, we identified 4 additional residues located in functionally-relevant regions (near the heme-binding region of the cytochrome b5 domain or the His boxes of the desaturase domain) that vary between these proteins (Supplementary Fig. 6).

**Figure 4.**
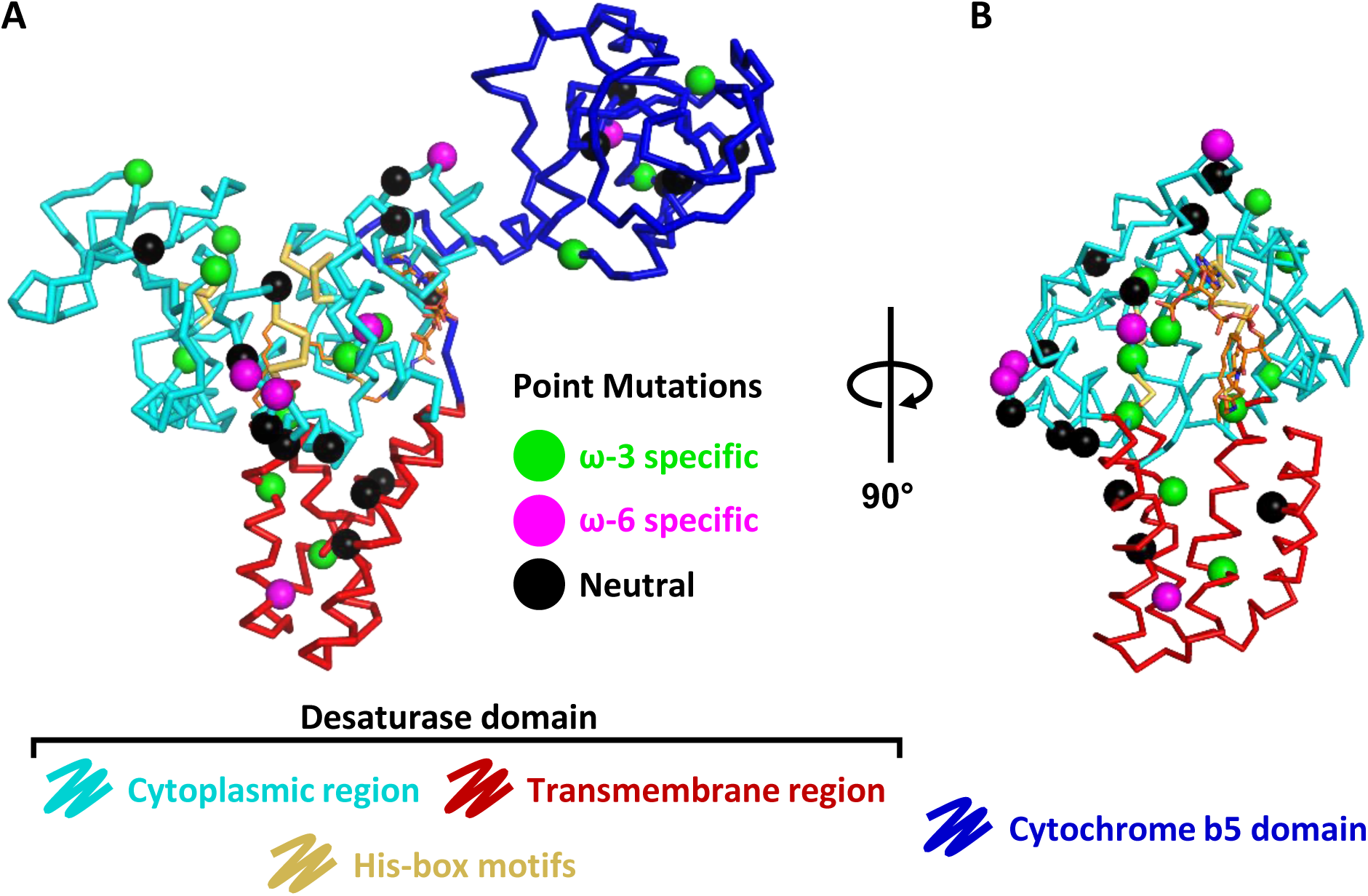
Position of specificity-altering MpΔ6des mutations. **A.** Ribbon representation of MpΔ6des with mutated residues identified as colored spheres as per their effect on enzyme substrate specificity. The conserved His-box motifs are shown in yellow and a [(Z)-octadec-9-enyl] (2R)-2,3-bis(oxidanyl)propanoate substrate analog is modeled bound to the enzyme active site based on analog-bound structures of MmΔ9des and HsΔ9des. Several mutations cluster near either the His-box motifs or the substrate binding site. Others, however, are located in distal positions such as the cytochrome b5 domain. **B.** A rotated view of the MpΔ6des ribbon with the cytochrome b5 domain hidden for clarity highlights the position of the substrate binding cleft.

**Figure 5.**
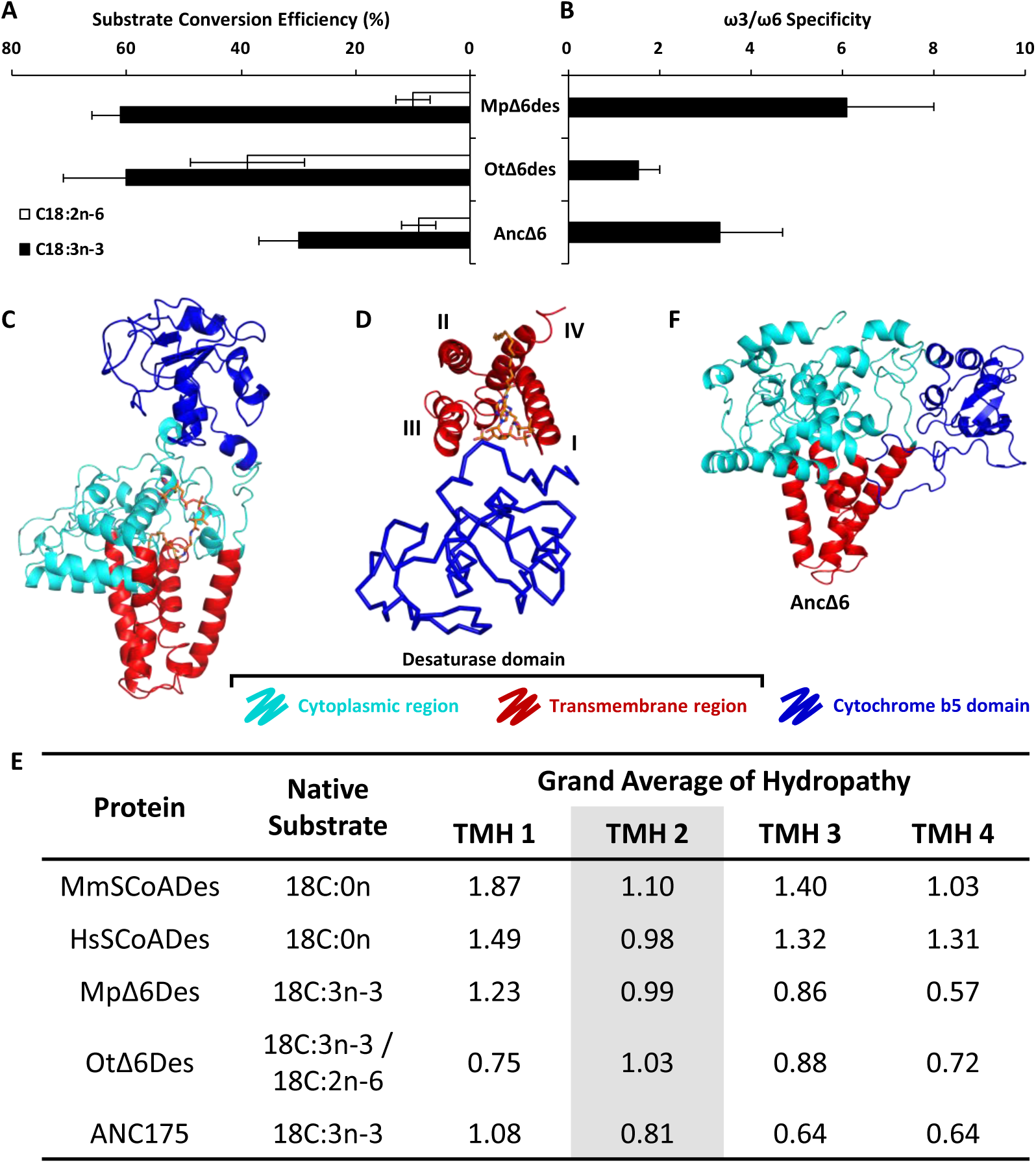
Structural insights into desaturase specificity from an ancestrally reconstructed Δ6-desaturase. **A.** Conversion efficiencies for both ω3 and ω6 substrates (ALA, black; and LA, white respectively) are shown for characterized Δ6-desaturases (n = 3 ± 1 S.D.) **B.** Relative ω3/ω6 specificities for characterized Δ6-desaturases are shown (n = 3 ± 1 S.D.), with AncΔ6 displaying strong ω3 specificity, albeit less so than MpΔ6des. **C.** The substrate binding pocket of Δ6-desaturases is formed by TMHs 1, 3, and 4 at the enzyme’s surface, with the cytochrome b5 domain positioned nearby due to its direct link to TMH 1. A [(Z)-octadec-9-enyl] (2R)-2,3-bis(oxidanyl)propanoate substrate analog molecule is modeled bound to the enzyme active site based on analog-bound structures of MmΔ9des and HsΔ9des. **D.** TMH numbering highlighting the orientation of helices around the substrate binding pocket. The b5 domain is shown as a ribbon to indicate its relative position. **E.** Grand average of hydropathy models based on primary sequence of the four TMHs shows a decreased hydropathy for TMHs 1, 3, and 4 in Δ6-desaturases relative to control Δ9 desaturases. **F.** The Phyre2 model of AncΔ6 demonstrates a potential interaction between the N-terminal loop of the cytochrome b5 domain in Δ6-desaturases and their substrate binding cleft.

Following identification of these putative specificity-linked positions, a series of MpΔ6des single mutants were generated by reverting the MpΔ6des amino acid at each position to the consensus amino acid. The resulting variants were expressed in yeast and compared to the wild-type MpΔ6des using a yeast competition assay to determine the effect of these mutations on enzyme activity against ω3 and ω6 substrates (C18:3n-3^Δ9,12,15^ α-linolenic acid (ALA) and C18:2n-3^Δ9,12^ linoleic acid (LA), respectively) (Fig. 3a, Supplementary Fig. 7, Supplementary Table 2). As Δ6-desaturases, including MpΔ6des, cannot be readily purified in an active form,^*55*^ whole-cell measured activity values (triplicate) were normalized against a wild-type MpΔ6des control included in every set of assays. The level of Δ6-desaturase expression was determined using Western blots and was comparable to wild-type MpΔ6des expression for all tested variants (Supplementary Fig. 7). Nonetheless, the ratio of the ω3/ω6 activities, which is unaffected by expression or activity levels, was used to determine the enzyme specificity (Fig. 3b, Supplementary Table 2). Out of 37 point mutations, 13 were shown to increase ω3 specificity by at least 1 SD, 6 were shown to increase ω6 specificity by at least 1 SD, and 18 were neutral (within 1 SD) in terms of their effect on substrate specificity.

Mapping mutation sites and effects onto our MpΔ6des model (Fig. 2a, 4a) shows one cluster of specificity-modifying mutations at positions 221-223 along with a dearth of nearby neutral mutations. This site lies in the enzyme’s substrate binding cleft near the triad of His box motifs and can therefore be easily rationalized as affecting specificity (Fig. 4b). This result is also consistent with previous studies of front-end desaturases, which demonstrated that mutations near His boxes could affect either substrate preference or the regioselectivity.^*56*^ The remaining mutations partition throughout the MpΔ6des structure displaying no clear pattern, highlighting the complex nature of the interactions governing desaturase substrate specificity. However, four specificity-linked mutations are found in the MpΔ6des cytochrome b5 domain, such as the V74I mutation that nearly doubles the enzyme ω3 to ω6 activity ratio relative to the wild type. Although the importance of the cytochrome b5 domain has been well documented in regard to electron transport during Δ6-desaturase catalysis, knowledge of specific interactions between the cytochrome b5 and desaturase domains remains elusive as the only structures of membrane-bound desaturases solved to date do not contain a cytochrome b5 domain. These results are thus, to our knowledge, the first evidence of a link between this domain and Δ6-desaturase substrate specificity. This is consistent with evidence that the front-end desaturases arose from an early fusion event with a cytochrome b5 protein, followed by the co-evolution of the cytochrome b5 domain with the desaturase,^*57*^ to form a cooperative substrate binding site.

In addition to the 37 single point mutants characterized above, we also generated three double mutants to better understand potential interacting positions throughout the MpΔ6des desaturase domain, each containing two of the three mutations amongst S201N, E222D, and A270L. These mutations were chosen as they are all found in OtΔ6des and individually increase ω6 activity. The resulting double mutants all demonstrate decreased ω3 specificity, with an additive effect observed when the A270L mutation is combined with either S201N or E222D. The effects of S201N and E222D on the other hand are not additive, with the S201N/E222D double mutant showing no significant changes in specificity compared to the S201N mutant alone. Introducing all three mutations into MpΔ6des however nonetheless leads to a further increased ω6 specificity compared to any double mutant, generating the least specific MpΔ6des variant tested and highlighting the complex interactions responsible for MpΔ6des specificity.

### Effects of substrate conformational restriction on Δ6-desaturase specificity

Along with identifying specificity-linked positions in Δ6-desaturases, our kinetics results (Fig. 3) show a distinct correlation between activity and specificity in these enzymes. Almost universally amongst tested mutations, those that further increase the ω3 specificity of MpΔ6des come at a cost to overall enzyme activity against both ω3 and ω6 substrates, while mutations that reduce ω3 specificity lead to increased ω6 activity rather than reduced ω3 activity. This correlation highlights the difficulty the enzyme faces in discriminating between ALA and LA. Both are acyl chains of the same length, differing only by a single double-bond that imparts additional rigidity and forces chain curvature into ALA compared to the more flexible LA. Strikingly, MpΔ6des catalysis is far more efficient on the more conformationally restricted ω3 substrate, suggesting that specificity in MpΔ6des and related desaturases is gated by substrate entropy and that conformational preorganization is crucial to efficient catalysis. Thus, further increases to ω3 specificity likely occur through further conformational restriction of the substrate via exclusion of conformational states that are accessible to ω6 substrates but are poorly populated in the more rigid ω3 substrates, in turn leading to overall decreased rates of catalysis. Likewise, a decreased ω3 specificity would occur through reduced conformational stringency, increasing ω6 activity without adversely affecting ω3 activity.

### Reconstruction of an ancestral Δ6-desaturase

Beyond the contributions of individual positions to specificity, the results of our consensus mutagenesis studies also hint at how substrate specificity evolved amongst members of the desaturase family. Although stemming from the larger set of sequences, mutations belonging to the consensus set of Δ4/Δ5/Δ6-desaturases are either neutral or enhance ω3 specificity (3 neutral and 7 ω3-specific mutations) suggesting an evolutionary bias towards ω3 substrates amongst front-end desaturases. Mutations from the set of algal Δ6-desaturases, however, do not bias towards either LA or ALA specificity (13 neutral, 5 ω3-specific, and 5 ω6-specific mutations). In addition, the four characterized members of the Δ6-desaturase clade display vastly different specificities, from ALA : LA conversion efficiencies of 63% : 5% for MpΔ6des to 71% : 73% for OtΔ6des (Supplementary Table 3). To address the question of whether the ω3 specificity of MpΔ6des, or the substrate promiscuity of OtΔ6des are recently diverged phenotypes, the substrate specificity of the putative common ancestor of the algal Δ6-desaturase family was tested. This ancestor, AncΔ6, was inferred from the algal Δ6 clade juncture using a phylogenetic tree (Fig. 1). The mean posterior probability of the resulting AncΔ6 sequence of 0.90 is comparable to that of other successfully reconstructed ancestral proteins.^*58*^ Notably, AncΔ6 exhibits greater full-length expression than the wild-type proteins, which is consistent with previous examples of proteins generated through ancestral sequence reconstruction (Supplementary Fig. 7).^*58*^ In terms of activity AncΔ6 displays comparable, if slightly lower activity to the wild-type proteins and is specific for ω3-fatty acids, albeit less specific than MpΔ6des (Fig. 5a,b). This suggests that the substrate promiscuity towards ω3-/ω6-substrates observed in OtΔ6des, could be a more recent adaptation in response to environmental changes. Notably, the sequence differences between AncΔ6 and MpΔ6des tend to cluster in the cytochrome b5 domain and the cytosolic loop of His Box I, implying a role for these regions in improved ω3 specificity. In contrast, the sequence differences between AncΔ6 and OtΔ6des tend to cluster in the cytochrome b5 domain and the region between His Box II and putative TMH3.

## DISCUSSION

Our results demonstrate that front-end desaturase substrate specificity is the result of subtle and complex interactions between residues throughout the enzyme. Notably, we show that the cytochrome b5 domain, previously thought to be involved primarily in electron transfer during catalysis,^*59*^ is also implicated in front-end desaturase substrate recognition. Considering the subtle structural difference between ω3- and ω6-substrates, these results suggest that the cytochrome b5 domain plays an additional role in refining the geometry of the substrate binding cavity and imply that it is physically part of the binding site, or modifies the shape/nature of the binding site via outer-shell interactions.^*60*^ Based on homology to known desaturase structures, as well as the cytosolic position of the desaturase domain N-terminus, we can putatively position the b5 domain of front-end desaturases in the cytosol near the cleft formed by TMHs 1, 3, and 4 (Fig. 5c,d). As this cleft contains the substrate binding pocket, the b5 domain would thus be poised to interact with both the substrate and the desaturase domain. Further examination of the TMH sequences in front-end desaturases also reveals that these three helices are less hydrophobic than those found in characterized desaturases lacking a fused b5 domain (Fig. 5e), while TMH 2, which does not contribute to forming this cleft, is similarly hydrophobic across both sets of desaturases. This increased hydrophilicity in front-end desaturases suggests a weaker membrane interaction and also explains the difficulty in accurately predicting TMH topology in MpΔ6des. In addition, the N-terminal portion of the cytochrome b5 domain in front-end desaturases shares little homology with cytochrome b5 proteins, instead consisting of a proline and charged amino acid-rich motif resembling intrinsically disordered regions in other proteins. (REF) The conformational flexibility of such a region coupled with weaker membrane interactions in the desaturase domain could allow for interactions between the b5 domain and the Δ6-desaturase substrate-binding pocket. Such a contact was observed in the AncΔ6 Phyre2 model, demonstrating potential complementarity of the b5 N-terminal tail and the TMH 1,3,4 cleft and a potential mechanism for substrate gating by the b5 domain (Fig. 5f).

In addition to examining their function, our reconstruction of an ancestral Δ6-desaturase also helps us to understand how these enzymes evolved. MpΔ6des was originally isolated from *M. pusilla* CCMP1545, which is identified from Atlantic Ocean (English Channel),^*61*^ and is one of the most ubiquitous marine algae strains.^*62*^ However, *M. pusilla* CCMP1545 is absent in the oligotrophic, high salt and high light, Mediterranean Sea, where *O. tauri* OTTH0595 is instead abundant.^*62, 63*^ Rather than geographical barriers driving divergence between species, differences between salt concentrations, light intensities and nutrient availabilities in some habitats might be an alternative selection pressure.^*64, 65*^ Increased levels of unsaturated fatty acids can maintain an adequate level of membrane fluidity and protect the photosynthesis machinery, thereby increases the salt tolerance of photosynthetic organisms.^*47, 66-68*^ Furthermore, excess photoassimilates are stored in the form of polyunsaturated fatty acids as a means to preventing oxidative stress under high light treatment.^*68-71*^ Ot6des activity produces comparatively elevated levels of ω6-fatty acids, which can then be incorporated into membranes and contribute to increased membrane ordering. Thus, it is expected that the promiscuity of this enzyme will result in a reduction in membrane fluidity given a similar level of LCPUFA incorporation. However, the higher temperatures in the Mediterranean Sea can mitigate these effects, promoting a higher rate of overall desaturation over specificity to increase fitness in a high-light environment. The colder environment of the Atlantic, on the other hand, requires organisms to develop higher membrane fluidity, hence promoting the evolution of more stringent Δ6-desaturases that display reduced rates of ω6-fatty acid production. Thus, we propose that the high salt concentration and high light of the Mediterranean Sea might have driven the divergence of the *O. tauri* Δ6-desaturase for high salt and light tolerance.

Although most enzyme engineering and evolution studies use enzymes that can be readily purified, easily assayed, or for which high-resolution structural information is available, desaturases present a more difficult system to study due to the lack of all three above factors. To circumvent these complicating factors, we used a combination of consensus mutagenesis, modelling, and ancestral sequence reconstruction to probe desaturase activity and specificity. Out of 37 consensus mutations found throughout the algal desaturase clades, all expressed and were active, and several encoded altered specificities that provided new insight into substrate specificity of these proteins. Several of the variants that we have produced could have value for biotechnology or as the basis for future biotech-based research, including variants with up to 8-fold increased ω-3 substrate specificity (albeit with loss of activity) and increased expression in yeast (AncΔ6) that could prove to be useful, depending on the rate-limiting factors in recombinant systems. Overall, our study has revealed new and useful insight into the structure-function relationship and evolution of these useful enzymes. This highlights the power of bioinformatics-inspired engineering approaches for generating small and manageable variant libraries for enzymes that are otherwise difficult to engineer, while also paving the way for the future engineering of industrially relevant LCPUFA desaturases.

## AUTHOR CONTRIBUTION

DL, CJ, TV, JP, and SS designed the experiments. DL performed all the experiments. AD, DL & CJ analyzed data with input from other authors. AD, DL & CJ wrote the manuscript with input from all authors.

## ACKNOWLEDGEMENTS

We thank Mr. Adam White, Dr. Pushkar Shrestha and Dr. Xue-Rong Zhou for their excellent technical assistance. We thank Professor Susan Howitt for providing the Western blot apparatus and nitrocellulose membranes used.

## REFERENCES

[1] Lee, J. H., O’Keefe, J. H., Lavie, C. J., and Harris, W. S. (2009) Omega-3 fatty acids: cardiovascular benefits, sources and sustainability, Nature reviews. Cardiology 6, 753–758.

[2] Chalon, S. (2006) Omega-3 fatty acids and monoamine neurotransmission, Prostaglandins, leukotrienes, and essential fatty acids 75, 259–269.

[3] Murphy, M. G. (1990) Dietary fatty acids and membrane protein function, The Journal of nutritional biochemistry 1, 68–79.

[4] Petrie, J. R., Shrestha, P., Mansour, M. P., Nichols, P. D., Liu, Q., and Singh, S. P. (2010) Metabolic engineering of omega-3 long-chain polyunsaturated fatty acids in plants using an acyl-CoA Delta6-desaturase with omega3-preference from the marine microalga Micromonas pusilla, Metab Eng 12, 233–240.

[5] Petrie, J. R., Shrestha, P., Zhou, X. R., Mansour, M. P., Liu, Q., Belide, S., Nichols, P. D., and Singh, S. P. (2012) Metabolic engineering plant seeds with fish oil-like levels of DHA, PloS one 7, e49165.

[6] Sayanova, O. V., Beaudoin, F., Michaelson, L. V., Shewry, P. R., and Napier, J. A. (2003) Identification of Primula fatty acid Delta(6)-desaturases with n-3 substrate preferences, Febs Lett 542, 100–104.

[7] Tang, X., Chen, H., Mei, T., Ge, C., Gu, Z., Zhang, H., Chen, Y. Q., and Chen, W. (2018) Characterization of an Omega-3 Desaturase From Phytophthora parasitica and Application for Eicosapentaenoic Acid Production in Mortierella alpina, Front Microbiol 9, e1878.

[8] Alonso, D. L., Garcia-Maroto, F., Rodriguez-Ruiz, J., Garrido, J. A., and Vilches, M. A. (2003) Evolution of the membrane-bound fatty acid desaturases, Biochem Syst Ecol 31, 1111–1124.

[9] Los, D. A., and Murata, N. (1998) Structure and expression of fatty acid desaturases, Biochim Biophys Acta 1394, 3–15.

[10] Vanhercke, T., Shrestha, P., Green, A. G., and Singh, S. P. (2011) Mechanistic and Structural Insights into the Regioselectivity of an Acyl-CoA Fatty Acid Desaturase via Directed Molecular Evolution, Journal of Biological Chemistry 286, 12860–12869.

[11] Na-Ranong, S., Laoteng, K., Kittakoop, P., Tanticharoen, M., and Cheevadhanarak, S. (2006) Targeted mutagenesis of a fatty acid Delta6-desaturase from Mucor rouxii: role of amino acid residues adjacent to histidine-rich motif II, Biochem Biophys Res Commun 339, 1029–1034.

[12] Sayanova, O., Beaudoin, F., Libisch, B., Castel, A., Shewry, P. R., and Napier, J. A. (2001) Mutagenesis and heterologous expression in yeast of a plant Delta6-fatty acid desaturase, J Exp Bot 52, 1581–1585.

[13] Shanklin, J., Whittle, E., and Fox, B. G. (1994) Eight histidine residues are catalytically essential in a membrane-associated iron enzyme, stearoyl-CoA desaturase, and are conserved in alkane hydroxylase and xylene monooxygenase, Biochemistry 33, 12787–12794.

[14] Avelange-Macherel, M. H., Macherel, D., Wada, H., and Murata, N. (1995) Site-directed mutagenesis of histidine residues in the delta 12 acyl-lipid desaturase of Synechocystis, Febs Lett 361, 111–114.

[15] Lindqvist, Y., Huang, W., Schneider, G., and Shanklin, J. (1996) Crystal structure of delta9 stearoyl-acyl carrier protein desaturase from castor seed and its relationship to other di-iron proteins, Embo J 15, 4081–4092.

[16] Dailey, H. A., and Strittmatter, P. (1979) Modification and identification of cytochrome b5 carboxyl groups involved in protein-protein interaction with cytochrome b5 reductase, J Biol Chem 254, 5388–5396.

[17] Hackett, C. S., and Strittmatter, P. (1984) Covalent cross-linking of the active sites of vesicle-bound cytochrome b5 and NADH-cytochrome b5 reductase, J Biol Chem 259, 3275–3282.

[18] Strittma, P., Spatz, L., Corcoran, D., Rogers, M. J., Setlow, B., and Redline, R. (1974) Purification and Properties of Rat-Liver Microsomal Stearyl Coenzyme-a Desaturase, P Natl Acad Sci USA 71, 4565–4569.

[19] Sayanova, O., Shewry, P. R., and Napier, J. A. (1999) Histidine-41 of the Cytochrome b5 Domain of the Borage Δ6 Fatty Acid Desaturase Is Essential for Enzyme Activity, Plant Physiol 121, 641–646.

[20] Zhou, X. R., Robert, S. S., Singh, S. P., and Green, A. G. (2006) Heterologous production of GLA and SDA by expression of an *Echium plantagineum* Δ6-desaturase gene, Plant Sci 172, 421–422.

[21] Napier, J. A., and Sayanova, O. (2005) The production of very-long-chain PUFA biosynthesis in transgenic plants: towards a sustainable source of fish oils, The Proceedings of the Nutrition Society 64, 387–393.

[22] Petrie, J. R., Shrestha, P., Belide, S., Kennedy, Y., Lester, G., Liu, Q., Divi, U. K., Mulder, R. J., Mansour, M. P., Nichols, P. D., and Singh, S. P. (2014) Metabolic engineering Camelina sativa with fish oil-like levels of DHA, PLoS One 9, e85061.

[23] Domergue, F., Abbadi, A., Zahringer, U., Moreau, H., and Heinz, E. (2005) In vivo characterization of the first acyl-CoA Delta6-desaturase from a member of the plant kingdom, the microalga Ostreococcus tauri, The Biochemical journal 389, 483–490.

[24] Diao, J., Song, X., Guo, T., Wang, F., Chen, L., and Zhang, W. (2019) Cellular engineering strategies toward sustainable omega-3 long chain polyunsaturated fatty acids production: State of the art and perspectives, Biotechnology Advances, e107497.

[25] Song, L. Y., Zhang, Y., Li, S. F., Hu, J., Yin, W. B., Chen, Y. H., Hao, S. T., Wang, B. L., Wang, R. R., and Hu, Z. M. (2013) Identification of the substrate recognition region in the Delta-fatty acid and Delta -sphingolipid desaturase by fusion mutagenesis, Planta.

[26] Meesapyodsuk, D., and Qiu, X. (2014) Structure Determinants for the Substrate Specificity of Acyl-CoA Delta9 Desaturases from a Marine Copepod, ACS chemical biology.

[27] Petrie, J. R., and Singh, S. P. (2011) Expanding the docosahexaenoic acid food web for sustainable production: engineering lower plant pathways into higher plants, AoB Plants 2011, plr011.

[28] Hoffmann, M., Wagner, M., Abbadi, A., Fulda, M., and Feussner, I. (2008) Metabolic engineering of omega3-very long chain polyunsaturated fatty acid production by an exclusively acyl-CoA-dependent pathway, The Journal of biological chemistry 283, 22352–22362.

[29] Hong, H., Datla, N., Reed, D. W., Covello, P. S., MacKenzie, S. L., and Qiu, X. (2002) High-level production of gamma-linolenic acid in Brassica juncea using a Delta 6 desaturase from Pythium irregulare, Plant Physiol 129, 354–362.

[30] Petrie, J. R., Liu, Q., Mackenzie, A. M., Shrestha, P., Mansour, M. P., Robert, S. S., Frampton, D. F., Blackburn, S. I., Nichols, P. D., and Singh, S. P. (2010) Isolation and characterisation of a high-efficiency desaturase and elongases from microalgae for transgenic LC-PUFA production, Mar Biotechnol (NY) 12, 430–438.

[31] Hitz, W. D., Carlson, T. J., Booth, J. R., Jr., Kinney, A. J., Stecca, K. L., and Yadav, N. S. (1994) Cloning of a higher-plant plastid omega-6 fatty acid desaturase cDNA and its expression in a cyanobacterium, Plant Physiol 105, 635–641.

[32] Winston, F., Dollard, C., and Ricupero-Hovasse, S. L. (1995) Construction of a set of convenient Saccharomyces cerevisiae strains that are isogenic to S288C, Yeast 11, 53–55.

[33] Pruitt, K. D., Tatsusova, T., and Maglott, D. R. (2007) NCBI reference sequences (RefSeq): a curated non-redundant sequence database of genomes, transcripts and proteins., Nucleic Acids Res 35, D61–65.

[34] Zhang, J., and Madden, T. L. (1997) PowerBLAST: a new network BLAST application for interactive or automated sequence analysis and annotation, Genome research 7, 649–656.

[35] Huang, Y., Niu, B., Gao, Y., Fu, L., and Li, W. (2010) CD-HIT Suite: a web server for clustering and comparing biological sequences, Bioinformatics 26, 680–682.

[36] Edgar, R. C. (2004) MUSCLE: multiple sequence alignment with high accuracy and high throughput, Nucleic Acids Res 32, 1792–1797.

[37] Tamura, K., Stecher, G., Peterson, D., Filipski, A., and Kumar, S. (2013) MEGA6: Molecular Evolutionary Genetics Analysis version 6.0, Molecular biology and evolution 30, 2725–2729.

[38] Yang, Z. (2007) PAML 4: phylogenetic analysis by maximum likelihood, Molecular biology and evolution 24, 1586–1591.

[39] Dobson, L., Remenyi, I., and Tusnady, G. E. (2015) CCTOP: A Consensus Constrained TOPology prediction web server, Nucleic Acids Res 43, W408–W412.

[40] Omasits, U., Ahrens, C. H., Muller, S., and Wollscheid, B. (2014) Protter: interactive protein feature visualization and integration with experimental proteomic data, Bioinformatics 30, 884–886.

[41] Kelley, L. A., Mezulis, S., Yates, C. M., Wass, M. N., and Sternberg, M. J. E. (2015) The Phyre2 web portal for protein modeling, prediction and analysis, Nat Protoc 10, 845–858.

[42] Bai, Y., McCoy, J. G., Levin, E. J., Sobrado, P., Rajashankar, K. R., Fox, B. G., and Zhou, M. (2015) X-ray structure of a mammalian stearoyl-CoA desaturase, Nature 524, 252–256.

[43] Wang, H., Klein, M. G., Zou, H., Lane, W., Snell, G., Levin, I., Li, K., and Sang, B. C. (2015) Crystal structure of human stearoyl-coenzyme A desaturase in complex with substrate, Nat Struct Mol Biol 22, 581–585.

[44] Zheng, L., Baumann, U., and Reymond, J. L. (2004) An efficient one-step site-directed and site-saturation mutagenesis protocol, Nucleic Acids Res 32, e115.

[45] Gibson, D. G., Young, L., Chuang, R. Y., Venter, J. C., Hutchison, C. A., 3rd, and Smith, H. O. (2009) Enzymatic assembly of DNA molecules up to several hundred kilobases, Nat Methods 6, 343–345.

[46] Tripodi, K. E., Buttigliero, L. V., Altabe, S. G., and Uttaro, A. D. (2006) Functional characterization of front-end desaturases from trypanosomatids depicts the first polyunsaturated fatty acid biosynthetic pathway from a parasitic protozoan, FEBS J 273, 271–280.

[47] Sakamoto, T., and Murata, N. (2002) Regulation of the desaturation of fatty acids and its role in tolerance to cold and salt stress, Current opinion in microbiology 5, 208–210.

[48] Svensk, E., Stahlman, M., Andersson, C. H., Johansson, M., Boren, J., and Pilon, M. (2013) PAQR-2 regulates fatty acid desaturation during cold adaptation in C. elegans, PLoS genetics 9, e1003801.

[49] Chinnusamy, V., Zhu, J., and Zhu, J. K. (2007) Cold stress regulation of gene expression in plants, Trends in plant science 12, 444–451.

[50] Michaud, M. R., and Denlinger, D. L. (2006) Oleic acid is elevated in cell membranes during rapid cold-hardening and pupal diapause in the flesh fly, Sarcophaga crassipalpis, J Insect Physiol 52, 1073–1082.

[51] Garcia-Maroto, F., Garrido-Cardenas, J. A., Vilches-Ferron, M. A., Manas-Fernandez, A., and Alonso, D. L. (2006) Evolution of ‘front-end’ desaturases in Echium (Boraginaceae), Biochem Syst Ecol 34, 327–337.

[52] Ai, H. W., Shaner, N. C., Cheng, Z., Tsien, R. Y., and Campbell, R. E. (2007) Exploration of new chromophore structures leads to the identification of improved blue fluorescent proteins, Biochemistry 46, 5904–5910.

[53] Diaz, A. R., Mansilla, M. C., Vila, A. J., and de Mendoza, D. (2002) Membrane topology of the acyl-lipid desaturase from Bacillus subtilis, Journal of Biological Chemistry 277, 48099–48106.

[54] Li, S. F., Song, L. Y., Zhang, G. J., Yin, W. B., Chen, Y. H., Wang, R. R., and Hu, Z. M. (2011) Newly identified essential amino acid residues affecting Delta8-sphingolipid desaturase activity revealed by site-directed mutagenesis, Biochem Biophys Res Commun 416, 165–171.

[55] Wilding, M., Nachtshatt, M., Speight, R., and Scott, C. (2017) An improved and general streamlined phylogenetic protocol applied to the fatty acid desaturase family, Mol Phylogenetics Evol 115, 50–57.

[56] Shi, H., Chen, H., Gu, Z., Song, Y., Zhang, H., Chen, W., and Chen, Y. Q. (2015) Molecular mechanism of substrate specificity for delta 6 desaturase from Mortierella alpina and Micromonas pusilla, J Lipid Res 56, 2309–2321.

[57] Gostincar, C., Turk, M., and Gunde-Cimerman, N. (2010) The Evolution of Fatty Acid Desaturases and Cytochrome b5 in Eukaryotes, J Membrane Biol 233, 63–72.

[58] Whitfield, J. H., Zhang, W. H., Herde, M. K., Clifton, B. E., Radziejewski, J., Janovjak, H., Henneberger, C., and Jackson, C. J. (2015) Construction of a robust and sensitive arginine biosensor through ancestral protein reconstruction, Protein Sci 24, 1412–1422.

[59] Napier, J. A., Michaelson, L. V., and Sayanova, O. (2003) The role of cytochrome b5 fusion desaturases in the synthesis of polyunsaturated fatty acids, Prostaglandins, leukotrienes, and essential fatty acids 68, 135–143.

[60] Hong, N.-S., Petrovic, D., Lee, R., Gryn’ova, G., Purg, M., Saunders, J., Bauer, P., Carr, P. D., Lin, C.-Y., Mabbitt, P. D., Zhang, W., Altamore, T., Easton, C., Coote, M. L., Kamerlin, S. C., and Jackson, C. J. (2018) The evolution of multiple active site configurations in a designed enzyme, Nat Commun 9, e3900.

[61] Vaulot, D., Le Gall, F., Marie, D., Guillou, L., and Partensky, F. (2004) The Roscoff Culture Collection (RCC): a collection dedicated to marine picoplankton, Nova Hedwigia 79, 49–70.

[62] Vaulot, D., Eikrem, W., Viprey, M., and Moreau, H. (2008) The diversity of small eukaryotic phytoplankton (< or =3 microm) in marine ecosystems, FEMS microbiology reviews 32, 795–820.

[63] Marie, D., Zhu, F., Balague, V., Ras, J., and Vaulot, D. (2006) Eukaryotic picoplankton communities of the Mediterranean Sea in summer assessed by molecular approaches (DGGE, TTGE, QPCR), FEMS microbiology ecology 55, 403–415.

[64] Rodriguez, F., Derelle, E., Guillou, L., Le Gall, F., Vaulot, D., and Moreau, H. (2005) Ecotype diversity in the marine picoeukaryote Ostreococcus (Chlorophyta, Prasinophyceae), Environmental microbiology 7, 853–859.

[65] Psarra, S., Tselepides, A., and Ignatiades, L. (2000) Primary productivity in the oligotrophic Cretan Sea (NE Mediterranean): seasonal and interannual variability, Prog Oceanogr 46, 187–204.

[66] Allakhverdiev, S. I., Kinoshita, M., Inaba, M., Suzuki, I., and Murata, N. (2001) Unsaturated fatty acids in membrane lipids protect the photosynthetic machinery against salt-induced damage in Synechococcus, Plant Physiol 125, 1842–1853.

[67] Zhang, J. T., Liu, H., Sun, J., Li, B., Zhu, Q., Chen, S. L., and Zhang, H. X. (2012) Arabidopsis Fatty Acid Desaturase FAD2 Is Required for Salt Tolerance during Seed Germination and Early Seedling Growth, Plos One 7.

[68] Klyachko-Gurvich, G. L., Tsoglin, L. N., Doucha, J., Kopetskii, J., Ryabykh, I. B. S., and Semenenko, V. E. (1999) Desaturation of fatty acids as an adaptive response to shifts in light intensity, Physiol Plantarum 107, 240–249.

[69] Solovchenko, A. E., Khozin-Goldberg, I., Didi-Cohen, S., Cohen, Z., and Merzlyak, M. N. (2008) Effects of light intensity and nitrogen starvation on growth, total fatty acids and arachidonic acid in the green microalga Parietochloris incisa, J Appl Phycol 20, 245–251.

[70] Mendoza, H., Martel, A., del Rio, M. J., and Reina, G. G. (1999) Oleic acid is the main fatty acid related with carotenogenesis in Dunaliella salina, J Appl Phycol 11, 15–19.

[71] Rabbani, S., Beyer, P., Lintig, J., Hugueney, P., and Kleinig, H. (1998) Induced beta-carotene synthesis driven by triacylglycerol deposition in the unicellular alga dunaliella bardawil, Plant Physiol 116, 1239–1248.

