## Supplementary Information for "Consensus mutagenesis and ancestral reconstruction provide insight into the substrate specificity and evolution of the front-end Δ6-desaturase family"

### Supplementary Figures

**The LCPUFA synthesis pathway**

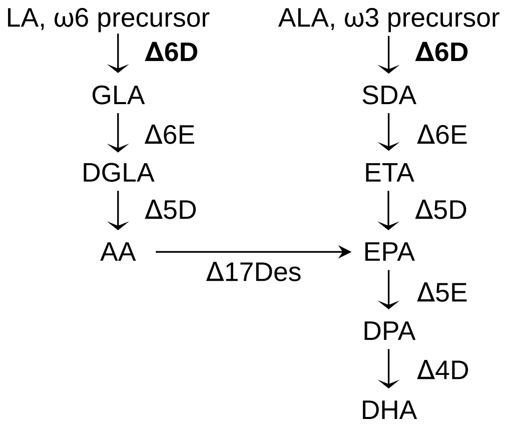


**Supplementary Figure 1. The marine algae LCPUFA synthesis pathway.** Using ω3 or ω6 substrates, marine algae are capable of producing LCPUFAs through a series of desaturation (Δ6D, Δ5D, and Δ4D) and elongation (Δ6E and Δ5E) steps. Δ6 desaturation is the rate-limiting step in the process. The fatty acid substrates and products along this pathway include: ALA (α-linolenic acid, $18:3^{\Delta 9,12,15}$), SDA (stearidonic acid, $18:4^{\Delta 6,9,12,15}$), ETA (eicosatetraenoic acid, $20:4^{\Delta 8,11,14,17}$), EPA (eicosapentaenoic acid, $20:5^{\Delta 5,8,11,14,17}$), DPA (docosapentaenoic acid, $22:5^{\Delta 7,10,13,16,19}$), DHA (docosahexaenoic acid, $22:6^{\Delta 4,7,10,13,16,19}$), LA (linoleic acid, $18:2^{\Delta 9,12}$), GLA (γ-linoleic acid, $18:3^{\Delta 6,9,12}$), DGLA (dihomo-γ-linoleic acid, $20:3^{\Delta 8,11,14}$), and AA (arachidonic acid,$20:3^{\Delta 5,8,11,14}$).

**% Identity**

| | | MpΔ6des | OtΔ6des | MmΔ9des | HsΔ9des | BsΔ5des |
|---|---|---|---|---|---|---|
| Algal | MpΔ6des | | 65.7 | 17.0 | 18.1 | 18.7 |
| | OtΔ6des | 65.7 | | 20.2 | 18.9 | 20.3 |
| Mammalian | MmΔ9des | 17.0 | 20.2 | | 85.1 | 15.6 |
| | HsΔ9des | 18.1 | 18.9 | 85.1 | | 15.5 |
| Bacterial | BsΔ5des | 18.7 | 20.3 | 15.6 | 15.5 | |

**Supplementary Figure 2. Sequence identity of characteristic algal Δ6-desaturases against desaturases of known topology.** Clustal Omega alignments of five desaturases (b5 domain not included for MpΔ6des and OtΔ6des) show low sequence identity for desaturases from different kingdoms. As no algal desaturase topology has been experimentally validated, sequence identity alone is insufficient to infer algal desaturase topology as identity levels are similar to both verified mammalian (4 TMH) and bacterial (6 TMH) topologies.

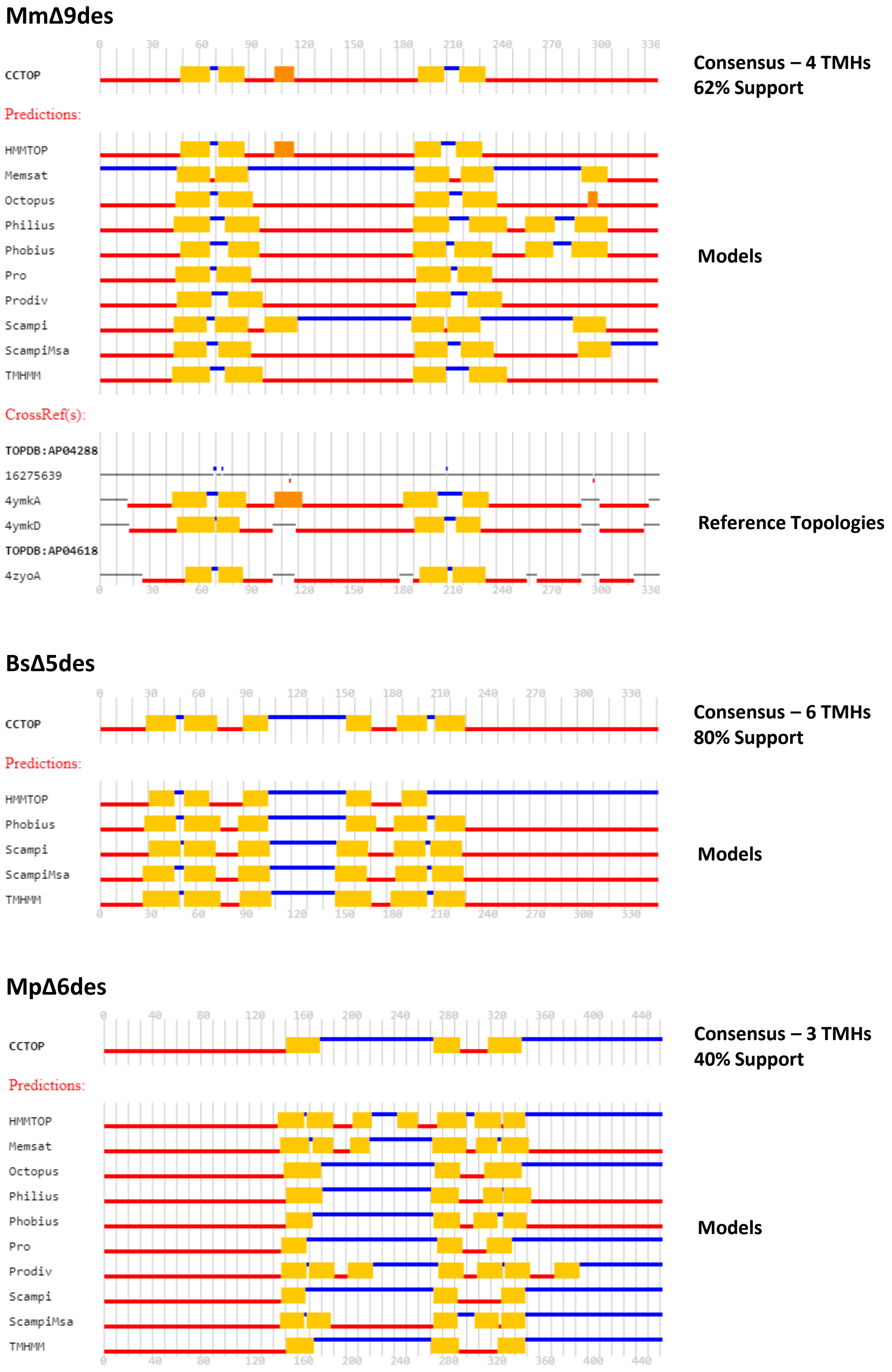


**Supplementary Figure 3. CCTOP topology predictions.** CCTOP topology predictions are shown for MpΔ6des and two control desaturases with known topologies (MmΔ6des – 4 TMH topology, and BsΔ5des – 6 TMH topology). The CCTOP prediction consists of a consensus from up to ten common topology modelling techniques, supported by available cross references from the TOPDB topology database. Listed support values correspond to the percentage of models (both predicted and referenced) that match the number of TMHs in the consensus. Residue numbering for MpΔ6des is offset by 130 residues relative to BsΔ5des and 106 residues relative to MmΔ9des due to the N-terminal b5 domain.

```
CCO20439.1          MEGAKKTSGRDSPTSSLAAHPGGRSSSSFFNNESHDEDDSSGGEFETEKKRKTTTREKDA  60
XP_003056992.1      -MCPPKTDGRSSP-------------------R-SPLTR-SKSSAEALDAKDASTAPVDL  38
XP_002502445.1      ------------------------------------MTR-GNKA---------KLDNSKL  14
CAQ30479.1          -MCPPKESTRKNA-------------------G-GPLTR-GK-------------LSADL  25
XP_003082578.1      -MCVETEN----------------------NDG-IPTVEIAFDGER---ERAEANVKLSA  33
DAA34893.1          -MCVETTEGTSRTMAN----ERTSSSSSLSEGG-TPTVTVGMGSEDAGKKTRNASVTAWT  54
BAK08911.1          ---------------------------------------MGRGGEG----------QVNS  11
ACD03117.1          ---------------------------------------MGKGSEG----------RSAA  11
XP_001562610.1      ------------------------------------------------------------  0
XP_003872262.1      ------------------------------------------------------------  0
AEI59766.1          ------------------------------------------------------------  0
XP_001681021.1      ------------------------------------------------------------  0
XP_001463326.1      ------------------------------------------------------------  0
XP_003858558.1      ------------------------------------------------------------  0
ABF58685.1          -------------M----------SS------LT----------LY----------RGPF  11
EGB03053.1          ---------------------------------------------M----------GKGA  5
EGD73438.1          ---------------------------------------------M----------GGGG  5
XP_001743517.1      -------------------------------------------MPS----------STKS  7
ABL96295.1          ---------------------------------MPPRDSYSYAAPP----------SAQL  17
AFD22890.1          -----MASKASSGI----------HS------STLPKEFHGATNDS----------RTDA  29
AEA72469.1          -MVAGKSGAAAHVT----------HS------STLPREYHGATNDS----------RSEA  33

                                      I                  I
CCO20439.1          KKMEPAQVAKTFERQWVKIGDLEYDVTDM--KHPGGSVIYYMLSN--VGADATEAFREFH  116
XP_003056992.1      KTLEPHELAATFETRWVRVEDVEYDVTNF--KHPGGSVIFYMLAN--TGADATEAFKEFH  94
XP_002502445.1      AKLEPHKLAQTFEQRWVRIDDVEYDVTNF--KHPGGSVIFYMLSN--TGADATEAFKEFH  70
CAQ30479.1          AKLEPHKLAQTFDTRWVRVGDVEYDVTNF--KHPGGSVIFYMLSN--TGADATEAFNEFH  81
XP_003082578.1      EKMEPAALAKTFARRYVVIEGVEYDVTDF--KHPGGTVIFYALSN--TGADATEAFKEFH  89
DAA34893.1          KELEPHAIAKTFERRYVTIEGVEYDVTDF--KHPGGSVIYYMLSN--TGADATEAFKEFH  110
BAK08911.1          VQVA---QGGAGTRKTILIEGEVYDVTNF--RHPGGSIIKFLTTDGTEAVDATNAFREFH  66
ACD03117.1          REMT-A-EANGDKRKTILIEGVLYDATNF--KHPGGSIINFLT-EGEAGVDATQAYREFH  66
XP_001562610.1      --MAPD-SALPHQPNEVLIDGVLYDCLNF--RHPGGSILKYYLGS----GDATETYRQFH  51
XP_003872262.1      --MTLD-SVLPHQPNEVLIDGVLYDCTDF--RHPGGGILRYYLGS----GDATETYRQFH  51
AEI59766.1          --MALD-NVRPHQPNEVLIDGVLYDCTDF--RHPGGSILKYYLGS----GDATETYQQFH  51
XP_001681021.1      --MALD-NVRPHQPNEVLIDGVLYDCTDF--RHPGGSILKYYLGS----GDATETYQQFH  51
XP_001463326.1      --MAPD-NARPHQRNEVLIDGVLYDCTDF--RHPGGSILKYYLGS----GDATETYQQFH  51
XP_003858558.1      --MAPD-NVRPHQRNEVLIDGVLYDCTDF--RHPGGSILKYYLGS----GDATETYQQFH  51
ABF58685.1          -------SRMVLPRQEICIDGRIYDVTEFINRHPGGKIILFQVG-----ADATDAFREFH  59
EGB03053.1          QQLTSP-AVKDAAAKDVLIHGQLYDVTNF--KHPGGSIVKFLTNN----GDATEAFVEFH  58
EGD73438.1          QQ-----QQPAKERKEVLIDGMLYDVTSF--RHPGGSIIKFLTNH----GDATDAFHQFH  54
XP_001743517.1      SA-----DQASSEVQEVLINGKLYDVSGF--KHPGGSVIKFLVNN----GDATEAFEEFH  56
ABL96295.1          HEVDTP---QEHDKKELVIGDRAYDVTNFVKRHPGGKIIAYQVG-----TDATDAYKQFH  69
AFD22890.1          PDLTIP---SLDPSKEMIIHGRVYDVSSFVKRHPGGSIIKFQLG-----ADATDAYNNFH  81
AEA72469.1          ADVTVS---SIDAEKEMIINGRVYDVSSFVKRHPGGSVIKFQLG-----ADASDAYNNFH  85
                                                    Heme-binding motif
                                                                                  E
CCO20439.1          YRSRKAEKMLKCLPQRKYDAKSHYSSKYNNNNSNYQPIDTSDSAMLKDFATFRLGLEKDG  176
XP_003056992.1      MRSLKAWKMLRALPSRPAEIKR---------------SESEDAPMLEDFARWRAELERDG  139
XP_002502445.1      MRSPKAWKMLKALPQRPAETPR---------------SADPDAPMLQDFARWRAELEKEG  115
CAQ30479.1          MRSPKAWKMLKALPNRPAETPR---------------SQDPDGPMLEDFAKWRAQLEKEG  126
XP_003082578.1      HRSRKARKALAALPSRPAKTA-----------------KVDDAEMLQDFAKWRKELERDG  132
DAA34893.1          YRSKKARKALAALPHKPVDAATR--------------EPIEDEAMLKDFAQWRKELEREG  156
BAK08911.1          CRSGKAEKYLKSLPKLGAPS-KMK--FDA-------KEQARRDAITRDYVKLREEMVAEG  116
ACD03117.1          QRSGKADKYLKSLPKLDASKVESR--FSA-------KEQARRDAMTRDYAAFREELVAEG  117
XP_001562610.1      MKLPRADKYLKLLPHRPAPPQHNVNV----------EEQKRLAKLSRDFKELQDACVEEG  101
XP_003872262.1      LKLPRADKYLKRLPNRPAPSQRSFNA----------DEHRRLAKLSRDFKALQDACVEEG  101
AEI59766.1          LKLPRADKYLKRLPNRPAPPQHSVNV----------DEQKRLEKLSRDFKALQDACVEEG  101
XP_001681021.1      LKLPRADKYLKRLPNRPAPPQHSVNV----------DEQKRLEKLSRDFKALQDACVEEG  101
XP_001463326.1      LKLPRADKYLKRLPNRPAPPQHGVDV----------DEQKRLAKLSRDFKALHDACVEEG  101
XP_003858558.1      LKLPRADKYLKRLPNRPAPPQHSVDV----------DEQKRLAKLSRDFKALHDACVEEG  101
ABF58685.1          AGSEKAEKILKTLPSRDDDGTFLPS---------------TQRSIMDDFKRLRDDLVSRG  104
EGB03053.1          GRSKKAQAMLKAMPKVAADAETMKARGYN-------GKEARSSALSVGYAKLRAEFEAEG  111
EGD73438.1          LRSERAQKLLKVLPKRPAPKDVIAERGGN-------GQEG----LAKAFAKLHSDLKAEG  103
XP_001743517.1      LRSKRAQYLLKSLPNRPAPKDVMVERGYN-------NREE----LTKDYAALRRQFKEEG  105
ABL96295.1          VRSAKADKMLKSLPSRPVHKGYSP----------------RRADLIADFQEFTKQLEAEG  113
AFD22890.1          MRSKKADKMLHSLPSRPAHADYAQ------------------DALSQDFEKLRLQLKEEG  123
AEA72469.1          VRSKKADKMLYSLPSRPAEAGYAQ------------------DDISRDFEKLRLELKEEG  127

                                                                   G     M
CCO20439.1          YFTPSLVHVAYRIVELMFTFALATYLMAKGYTTL---SVITYGAFFGARCGWVQHEGGHN  233
XP_003056992.1      FFKPSITHVAYRLLELLATFALGTALMYAGYPII---ASVVYGAFFGARCGWVQHEGGHN  196
XP_002502445.1      FFEPSRLHLAYRCLELCATFALGTFLMYIGRPLL---ASIVYGAFFGARCGWVQHEGGHN  172
CAQ30479.1          FFKPSIAHVAYRIAELAAMFALGCYIMSLGYPVV---ASIVFGAFFGARCGWVQHEGGHN  183
XP_003082578.1      FFKPSPAHVAYRFAELAAMYALGTYLMYARYVVS---SVLVYACFFGARCGWVQHEGGHS  189
DAA34893.1          FFKPSPAHVAYRFAELAAMFALGTALMHARWHVA---SVIVYSCFFGARCGWVQHEGGHN  213
BAK08911.1          LFKPAPLHIVYRFAEIAALFAASFYLFSMRGNVFATLAAIAVGGIAQGRCGWLMHECGHF  176
ACD03117.1          YFDPSIPHMIYRVVEIVALFALSFWLMSKASPTS-LVLGVVMNGIAQGRCGWVMHEMGHG  176
XP_001562610.1      LFDPSWPHIFYRFSELILMHVVGFYILFRLSLLWP--IALVILGVAEGRCGWWMHEAGHY  159
XP_003872262.1      LFNASWPHIVYRFSELILMHAIGLYMLFRLPMLWP--IALLILGVAEGRCGWWMHEAGHY  159
AEI59766.1          LFNASWPHIVYRFSELILMHAIGLYMLFRLPILWP--VALVILGVAEGRCGWWMHEAGHY  159
XP_001681021.1      LFNASWPHIVYRFSELILMHAIGLYMLFRLPILWP--VALVILGVAEGRCGWWMHEAGHY  159
XP_001463326.1      LFNASWPHIVYRFSELILMHAIGLYMLFHLPMLWP--IALVILGVAEGRCGWWMHEAGHY  159
XP_003858558.1      LFNASWPHIVYRFSELILMHAIGLYMLFRLPMLWP--IALVILGVAEGRCGWWMHEAGHY  159
ABF58685.1          VFKPSVMHVVYRCLEVVALYLIGFYLALCTSNVY---VGCAVLGVAQGRAGWLMHEGGHH  161
EGB03053.1          RFDANASEIAYRIGEVVLMHLVGAYLIMATDHFF---KGLLCLGIVSGRCGWLMHEGGHY  168
EGD73438.1          YFDPSMAEVAYRTLELVAIHALGAYFLMTATNVFSVIAGILCLAMGQGRCGWLMHEGGHY  163
XP_001743517.1      LFEPSRGEIALRFAEIIGMHVLGAYLVWQGGL--FTALGVVMLAIVQGRCGWLMHEGGHY  163
ABL96295.1          MFEPSLPHVAYRLAEVIAMHVAGAALIWHGYTF----AGIAMLGVVQGRCGWLMHEGGHY  169
AFD22890.1          YFEPNLRHVAYRIIEVIAMYWAGIQLIWSGYWF----LGAIVAGLAQGRCGWLQHEGGHY  179
AEA72469.1          YFEPNLVHVSYRCVEVLAMYWAGVQLIWSGYWF----LGAIVAGIAQGRCGWLQHEGGHY  183
                              TMH I                  TMH II               His-box I
```

```
                                        A W
CCO20439.1        SLTGNIWIDKRIQACLMGFGLGTSGDMWNVMHNKHHATPQKIRHDMDLDTTPAVAFFNTA  293
XP_003056992.1    SLTGSVYVDKRLQAMTCGFGLSTSGEMWNQMHNKHHATPQKVRHDMDLDTTPAVAFFNTA  256
XP_002502445.1    SLTGSIWWDKRIQAATCGFGLSTSGDMWNQMHNKHHATPQKVRHDMDLDTTPAVAFFDTA  232
CAQ30479.1        SLTGNIWLDKRIQAATCGFGLSTSGDMWNQMHNKHHATPQKVRHDMDLDTTPAVAFFKTA  243
XP_003082578.1    SLTGNIWWDKRIQAFTAGFGLAGSGDMWNSMHNKHHATPQKVRHDMDLDTTPAVAFFNTA  249
DAA34893.1        SLTGNIWWDKRIQAFAAGFGLASSGDMWNNMHNKHHATPQKVRHDMDLDTTPTVAFFNSA  273
BAK08911.1        SMTGYIPLDVRLQELVYGVGCSMSASWWRVQHSKHHATPQKLKHDVDLDTLPLVAFNEKI  236
ACD03117.1        SFTGVIWLDDRMCEFFYGVGCGMSGHYWKNQHSKHHAAPNRLEHDVDLNTLPLVAFNERV  236
XP_001562610.1    SVTGIPWLDIKIQEVLYGLGDGMSASWWRSQHNKHHATPQKHHHDVDLETLPLVAFNTII  219
XP_003872262.1    SVTGIPWLDIKIQEVLYGLGDGMSASWWRSQHNKHHATPQKHRHDVDLETLPLVAFNKVI  219
AEI59766.1        SVTGIPWLDIKIQEVLYGLGDGMSASWWRSQHNKHHATPQKHRHDVDLETLPLVAFNKII  219
XP_001681021.1    SVTGIPWLDIKIQEVLYGLGDGMSASWWRSQHNKHHATPQKHRHDVDLETLPLVAFNKII  219
XP_001463326.1    SVTGIPWLDIKIQEVLYGLGDGMSASWWRSQHNKHHATPQKHRHDVDLETLPLVAFNKII  219
XP_003858558.1    SVTGIPWLDIKIQEVLYGLGDGMSASWWRSQHNKHHATPQKHRHDVDLETLPLVAFNKII  219
ABF58685.1        SLTGNWKVDQFLQELFFGIGCGMSAAWWRNAHNKHHAAPQHLGKDVDLETLPLVAFNKAV  221
EGB03053.1        SLTGVIKTDRFLQEAIYGVGCGMSAAWWRNQHNKHHATPQKLQHDVDLDTLPLVAFHAQV  228
EGD73438.1        SMTGNVAIDKAFQIVLYGVGCGMSAGWWRSQHNRHHATPQKLQHDVDLETLPLVAFNTRI  223
XP_001743517.1    SITGRVMLDRGLQIALYGIGCGMSAGWWRSQHNRHHATPQKLQHDVDLETLPLIAFNSVI  223
ABL96295.1        SLTGNIAFDRAIQVACYGLGCGMSGAWWRNQHNKHHATPQKLQHDVDLDTLPLVAFHERI  229
AFD22890.1        SLTGNIKIDRHLQMAIYGLGCGMSGCYWRNQHNKHHATPQKLGTDPDLQTMPLVAFHKII  239
AEA72469.1        SLTGNIKIDRHLQMAIYGLGCGMSGCYWRNQHNKHHATPQKLGTDPDLQTMPLVAFHKIV  243
                                                His-box II
                                          V
CCO20439.1        VEENRDRGFSRLWSRFQAWTFVPITSGVFVMAFWLYVLHPSKVFKKKNWEEAFWMLSSHV  353
XP_003056992.1    VEDNRPRGFSRAWARLQAWTFVPVTSGLLVQAFWIYVLHPRQVLRKKNYEEASWMLVSHV  316
XP_002502445.1    VEDNRPRGFSKTWARAQAWTFVPITSGVLVQMFWIYVLHPRQVLRKKNYEEASWMFLSHV  292
CAQ30479.1        VEDNRPRGFSRAWSRAQAWTFVPVTSGLLVQMFWIYVLHPRQVARKKNYEEASWMILSHV  303
XP_003082578.1    VEDNRPRGFSKYWLRLQAWTFIPVTSGLVL-LFWMFFLHPSKALKGGKYEELVWMLAAHV  308
DAA34893.1        VEENRPRGFSKLWLRLQAWTFVPVTSGMVL-FFWMFVLHPRNALRRKSFEEAAWMFSAHV  332
BAK08911.1        AAKVRPGSFQAKWLSAQAYIFAPVSCFLVG-LFWTLFLHPRHMLRTSHFAEMAAVAVRVV  295
ACD03117.1        VRKVKPGSLLALWLRVQAYLFAPVSCLLIG-LGWTLYLHPRYMLRTKRHMEFVWIFARYI  295
XP_001562610.1    ARRGRRSVNIRRWISLQRYLFAPVTCSLVA-LYWQLFLHVRHAMRTRRYTEGAAILCRWI  278
XP_003872262.1    ARRGKRNANIRGWISLQMYLFGPVTCSLVA-LYWQLFLHVRHAMRTQRYTEGAAILCRWI  278
AEI59766.1        ARRGKRNASIRRWISLQMFLFGPVTCSLVA-LYWQLFLHVRHAMRTQRYTEGSAILCRWI  278
XP_001681021.1    ARRGKRNASIRRWISLQMFLFGPVTCSLVA-LYWQLFLHVRHAMRTQRYTEGSAILCRWI  278
XP_001463326.1    ALRGKRNANIRRWISLQMFLFGPVTCSLVA-LYWQLFLHVRHAMRTQRYTEGVAILCRWI  278
XP_003858558.1    ALRGKRIANIRRWISLQMFLFGPVTCSLVA-LYWQLFLHVRHAMRTQRYTEGVAILCRWI  278
ABF58685.1        ---LR-GRLPSVWIRSQAVCFAPISTLLVS-FFWQFYLHPRHIIRTGRRMESFWLLVRYL  276
EGB03053.1        AAKAK-GALMKQWLKLQCYLFIPLSCLLVA-SGWQLYLHPRHAHRTKRGKELAFMGLRYV  286
EGD73438.1        AERTK-SPAVRAWLSMQHILFIPVSCLLVA-LGWQLFLHPRYMLRTKKKFEFMTLFVRYY  281
XP_001743517.1    AAKAR-NPLVKLWLRAQAFCFIPLSCLLVA-LGWQIYLHPRYMIRTKKRFELLTLAVRYL  281
ABL96295.1        AAKVK-SPAMKAWLSMQAKLFAPVTTLLVA-LGWQLYLHPRHMLRTKHYDELAMLGIRYG  287
AFD22890.1        AGQAK-GAKGKAWLAWQAPLFFGGIICSLVSFGWQFVLHPKHALRVQNHLELAYMALRYV  298
AEA72469.1        GAKAR--GKGKAWLAWQAPLFFGGIICSLVSFGWQFVLHPNHALRVHNHLELAYMGLRYV  301
                                                             TMH III
                                                    I
CCO20439.1        VRASLIQFVHPSNVSFAHAYGLYALSQWIAGMYLFAHFSTSHTHTKVVEFKDHPSWVRYA  413
XP_003056992.1    VRTAVIKLAT--GYSWPVAYWWFTFGNWIAYMYLFAHFSTSHTHLPVVPSDKHLSWVNYA  374
XP_002502445.1    VRTAVIKLAL--GCGTAEAYGWFWVGNWIAYMYLFAHFSTSHTHLDVVPSDKHISWVNYA  350
CAQ30479.1        LRTATIKYAG--GYSWPVAYLWFSFGNWIAYMYLFAHFSTSHTHLEVVPSDKHISWVNYA  361
XP_003082578.1    IRTWTIKAVT--GFTAMQSYGLFLATSWVSGCYLFAHFSTSHTHLDVVPADEHLSWVRYA  366
DAA34893.1        IRTAVIKAVT--GYSWIASYGLFAATMWASGCYLFAHFSTSHTHLDVVPSDKHLSWVRYA  390
BAK08911.1        G-WAALMHSF--GYSGSDSFGLYMATFGFGCTYIFTNFAVSHTHLDVTEPDEFLHWVEYA  352
ACD03117.1        G-WFSLMGAL--GYSPGTSVGMYLCSFGPGCIYIFLQFAVSHTHLPVTNPEDQLHWLEYA  352
XP_001562610.1    A-VGVICHKL--QVSFWQGLGGVLFSQAFGAAYIFISFALNHTHLPMLPEDQHAHFVEYA  335
XP_003872262.1    A-VGVICHQL--QVSFWQGLGGVLFSQAFSAAYIFINFALNHSHLPMLPEDEHAHFVEYA  335
AEI59766.1        V-VGVICHQL--QVSFWQGLGGVLFSQAFSAAYIFINFALSHSHLPMLPEDEHAHFVEYA  335
XP_001681021.1    V-VGVICHQL--QVSFWQGLGGVLFSQAFSAAYIFINFALNHSHLPMLPEDEHAHFVEYA  335
XP_001463326.1    A-VGVICHKL--QVSFWQGLGGVLFSQAFSAAYIFINFALNHSHLPMLPEDEHAHFVEYA  335
XP_003858558.1    A-VGVICHKL--QVSFWQGLGGVLFSQAFSAAYIFINFALNHSHLPMLPEDEHAHFVEYA  335
ABF58685.1        VIVY-LGFS----YGLVSVLLCYIASVHVGGMYIFVHFALSHTHLPVINQHGRANWLEYA  331
EGB03053.1        LLFGVVFKD----LTWLQALGYYNLYNQIAASYIFTNFSLSHTHLPVSQPDDYLHWVEYA  342
EGD73438.1        LLFGVVLQG----YTWPAAIAIYLLYNGLSASYIFTNFALSHTHLPVTNPDEYLHWVEYA  337
XP_001743517.1    VIFGVVLRD----FTWAQAIGLYILYDMIGAAYIFTNFALSHTHLPVSQADEYLHWVEYA  337
ABL96295.1        L-VGYLAAN----YGAGYVLACYLLYVQLGAMYIFCNFAVSHTHLPVVEPNEHATWVEYA  342
AFD22890.1        L-WHCAFGY----LGLLGSLRLYAFYVAVGGTYIFTNFAVSHTHKDVVPPTKHISWCLYS  353
AEA72469.1        L-WHLAFGH----LGLLSSLRLYAFYVAVGGTYIFTNFAVSHTHKDVVPPTKHISWALYS  356
                                    TMH IV
                        N
CCO20439.1        VEHTVDISPDRAYVNWLMGYLNCQVIHHLFPDMPQFRQPE-VSKKFELFARKWGLEYTVM  472
XP_003056992.1    VDHTVDIDPSRGYVNWLMGYLNCQVIHHLFPDMPQFRQPE-VSRRFVPFAKKWGLNYKVL  433
XP_002502445.1    VDHTVDINPRNSIVNWLMGYLNCQVIHHLFPDMPQFRQPE-VSRRFVAFAEKWNLNYKVL  409
CAQ30479.1        VDHTVDIDPSKGYVNWLMGYLNCQVIHHLFPDMPQFRQPE-VSRRFVAFAKKWNLNYKVL  420
XP_003082578.1    VDHTIDIDPSQGWVNWLMGYLNCQVIHHLFPSMPQFRQPE-VSRRFVAFAKKWNLNYKVM  425
DAA34893.1        VDHTIDINPNNSVVNWLMGYLNCQVIHHLFPDMPQFRQPE-VSRRFVPFAKKWNLNYKVL  449
BAK08911.1        ALHTTNVSNDSWFITWWMSYLNFQIEHHLFPSLPQLNAPR-VAPRVRALFEKHGMAYDER  411
ACD03117.1        ADHTVNISTKSWLVTWWMSNLNFQIEHHLFPTAPQFRFKE-ISPRVEALFKRHNLPYYDL  411
XP_001562610.1    AVYTMNVTP-SWLVTWFMGYLNYQVEHHLFPSMPQFRFVQ-LAPRVRRLFEENGLTYDSR  393
XP_003872262.1    AIYTMNVTP-SWFVTWFMGYLNYQVEHHLFPTMPQFRFPQ-LAPRVRKLFEENGLKYDSR  393
AEI59766.1        AIYTMNVTP-SWFVTWFMGYLNYQVEHHLFPTMPQFRFVQ-QAPRVRKLFEENGLKYDSR  393
XP_001681021.1    AIYTMNVTP-SWFVTWFMGYLNYQVEHHLFPTMPQFRFVQ-LAPRVRKLFEENGLKYDSR  393
XP_001463326.1    AIYTMNVTP-SWFVTWFMGYLNYQVEHHLFPTMPQFRFVQ-LAPRVRKLFEENGLKYDSR  393
XP_003858558.1    AIYTMNVTP-SWFVTWFMGYLNYQVEHHLFPTMPQFRFVQ-LAPRVRKLFEENGLKYDSR  393
ABF58685.1        SKHTVNVSTNNYFVTWLMSYLNYQIEHHLFPSCPQFRFPGYVSMRVREFFHKHGLKYNEV  391
EGB03053.1        ALHTTNISP-GLVCNWWMANLNFQIEHHLFPSMPQFQHQH-ISPRVKAFFEAHGLHYDVR  400
EGD73438.1        SKHTTNITS-SALCDWWMAYLNYQIEHHLFPSMPQFRHPK-IAGRVRKLFEDHGLVYDVR  395
XP_001743517.1    AKHTTNIAG-TPLCNWWMGYLNFQIEHHLFPCMPQFRHPQ-ISPRVKALFEKHGLPYDVR  395
ABL96295.1        ANHTTNCSP-SWWCDWWMSYLNYQIEHHLYPSMPQFRHPK-IAPRVKQLFEKHGLHYDVR  400
AFD22890.1        ANHTTNCTN-SPFVNWWMAYLNFQIEHHLFPSMPQYNHPT-IAPRVRALFEKHGVEYDVR  411
AEA72469.1        ANHTTNCSD-SPFVNWWMAYLNFQIEHHLFPSMPQYNHPK-IAPRVRALFEKHGVEYDVR  414
                                         His-box III
```

```
CCO20439.1          TYGEAWKATFKNLNDVGKHYYEEGRVKDKERSSEENAKTKKIK       515
XP_003056992.1      SYYGAWKATFSNLDKVGQHYYVNGKAEKAH-------------       463
XP_002502445.1      TYYGAWKATFSNLDRVGQHYYVNGKAKAH--------------       438
CAQ30479.1          TYYGAWKATFTNLDTVGQHYYKHGKAHAH--------------       449
XP_003082578.1      TYAGAWKATLGNLDNVGKHYYVHGQHSGKTA------------       456
DAA34893.1          TYYGAWKATFGNLNDVGKHYYVHGSQRVKSKSA----------       482
BAK08911.1          PYPTALGDTFANLHAVGQNAGQAAAKAA---------------       439
ACD03117.1          PYTSAVSTTFANLYSVGHSVGADTKKQD---------------       439
XP_001562610.1      PYAKSLQTTFKNLSDVAEFIVAGK-------------------       417
XP_003872262.1      PYMESLQKTFKNLGDVAAFIVAEN-------------------       417
AEI59766.1          PYMESLQKTFKNLGDVAEFIVAGN-------------------       417
XP_001681021.1      PYMESLQKTFKNLGDVAEFIVAGN-------------------       417
XP_001463326.1      PYMKSLQKTFKNLGDVAELIVAGN-------------------       417
XP_003858558.1      PYMKSLQKTFKNLGDVAEFIVAGN-------------------       417
ABF58685.1          GYLHALNLTFSNLAAVAIVE-----------------------       411
EGB03053.1          PYFSCLKQTLENLHDVGHSTDAVKKD-----------------       426
EGD73438.1          DYFACLDDTLSNLARVGNPDKAK--------------------       418
XP_001743517.1      GYWSCLGDTLSNMHQVGNPHAKAA-------------------       419
ABL96295.1          GYFEAMADTFANLDNVAHAPEKKMQ------------------       425
AFD22890.1          PYLECFRVTYVNLLAVGNPQHSYHTHTH---------------       439
AEA72469.1          PYLECFRVTYVNLLAVGNPEHSYHEHTH---------------       442
```

**Supplementary Figure 4. The alignment of algae Δ4/5/6-desaturases.** The consensus positions are shown in red with the consensus residues identified above the alignment. Functional motifs are highlighted in yellow and identified below the alignment. Putative MpΔ6des TMHs are underlined and identified below the alignment.

```
BpΔ6des                    MEGAKKTSGRDSPTSSLAAHPGGRSSSSFFNNESHDEDDSSGGEFETEKKRKTTTREKDA 60
OtΔ6des                    -MCVETENND-----------------------GIPTVEIAFDGE---RERAEANVKLSA 33
OlΔ6des                    -MCVETTEGTSRTMANERT----SSSSSLSE-GGTPTVTVGMGSEDAGKKTRNASVTAWT 54
MpΔ6des_CCMP1545           -MCPPKTDGRSSPR------------SPLTR-S--------KSSAEALDAKDASTAPVDL 38
AncΔ6                      -MCPPKTDGRSSPR------------SPLTR-S--------KSSAEALDAKDASTAPVDL 38
MpΔ6des_RCC299             ----------------------------MTR-G--------NKA---------KLDNSKL 14
MsΔ6des                    -MCPPKESTRKNAG------------GPLTR-G--------K-------------LSADL 25

                                        R             D             S              Y
BpΔ6des                    KKMEPAQVAKTFERQWVKIGDLEYDVTDMKHPGGSVIYYMLSNVGADATEAFREFHYRSR 120
OtΔ6des                    EKMEPAALAKTFARRYVVIEGVEYDVTDFKHPGGTVIFYALSNTGADATEAFKEFHHRSR 93
OlΔ6des                    KELEPHAIAKTFERRYVTIEGVEYDVTDFKHPGGSVIYYMLSNTGADATEAFKEFHYRSK 114
MpΔ6des_CCMP1545           KTLEPHELAATFETRWVRVEDVEYDVTNFKHPGGSVIFYMLANTGADATEAFKEFHMRSL 98
AncΔ6                      KTLEPHELAATFETRWVRIEGVEYDVTDFKHPGGSVIFYMLSNTGADATEAFKEFHYRSR 98
MpΔ6des_RCC299             AKLEPHKLAQTFEQRWVRIDDVEYDVTNFKHPGGSVIFYMLSNTGADATEAFKEFHMRSP 74
MsΔ6des                    AKLEPHKLAQTFDTRWVRVGDVEYDVTNFKHPGGSVIFYMLSNTGADATEAFNEFHMRSP 85
                                                         Heme-binding Motif
                                 A
BpΔ6des                    KAEKMLKCLPQRKYDAKSHYSSKYNNNNSNYQPIDTSDSAMLKDFATFRLGLEKDGYFTP 180
OtΔ6des                    KARKALAALPSRPAKTA-----------------KVDDAEMLQDFAKWRKELERDGFFKP 136
OlΔ6des                    KARKALAALPHKPVDAATR--------------EPIEDEAMLKDFAQWRKELEREGFFKP 160
MpΔ6des_CCMP1545           KAWKMLRALPSRPAEIK-R--------------SESEDAPMLEDFARWRAELERDGFFKP 143
AncΔ6                      KAWKMLKALPQRPADYK-R--------------AETEDAAMLKDFAKWRKELERDGFFKP 143
MpΔ6des_RCC299             KAWKMLKALPQRPAETP-R--------------SADPDAPMLQDFARWRAELEKEGFFEP 119
MsΔ6des                    KAWKMLKALPNRPAETP-R--------------SQDPDGPMLEDFAKWRAQLEKEGFFKP 130

                            P            M                 I                        NIW
BpΔ6des                    SLVHVAYRIVELMFTFALATYLMAKGYTTLSVITYGAFFGARCGWVQHEGGHNSLTGNIW 240
OtΔ6des                    SPAHVAYRFAELAAMYALGTYLMYARYVVSSVLVYACFFGARCGWVQHEGGHSSLTGNIW 196
OlΔ6des                    SPAHVAYRFAELAAMFALGTALMHARWHVASVIVYSCFFGARCGWVQHEGGHNSLTGNIW 220
MpΔ6des_CCMP1545           SITHVAYRLLELLATFALGTALMYAGYPIIASVVYGAFFGARCGWVQHEGGHNSLTGSVY 203
AncΔ6                      SLAHVAYRFVELAAMFALGTYLMYAGYWFSSVIVYGAFFGARCGWVQHEGGHNSLTGNIW 203
MpΔ6des_RCC299             SRLHLAYRCLELCATFALGTFLMYIGRPLLASIVYGAFFGARCGWVQHEGGHNSLTGSIW 179
MsΔ6des                    SIAHVAYRIAELAAMFALGCYIMSLGYPVVASIVFGAFFGARCGWVQHEGGHNSLTGNIW 190
                                 TMH I                  TMH II             His-box I
                           W   I  F          D
BpΔ6des                    IDKRIQACLMGFGLGTSGDMWNVMHNKHHATPQKIRHDMDLDTTPAVAFFNTAVEENRDR 300
OtΔ6des                    WDKRIQAFTAGFGLAGSGDMWNSMHNKHHATPQKVRHDMDLDTTPAVAFFNTAVEDNRPR 256
OlΔ6des                    WDKRIQAFAAGFGLASSGDMWNNMHNKHHATPQKVRHDMDLDTTPTVAFFNSAVEENRPR 280
MpΔ6des_CCMP1545           VDKRLQAMTCGFGLSTSGEMWNQMHNKHHATPQKVRHDMDLDTTPAVAFFNTAVEDNRPR 263
AncΔ6                      WDKRIQAFTAGFGLGTSGDMWNNMHNKHHATPQKVRHDMDLDTTPLVAFFNTAVEDNRPR 263
MpΔ6des_RCC299             WDKRIQAATCGFGLSTSGDMWNQMHNKHHATPQKVRHDMDLDTTPAVAFFDTAVEDNRPR 239
MsΔ6des                    LDKRIQAATCGFGLSTSGDMWNQMHNKHHATPQKVRHDMDLDTTPAVAFFKTAVEDNRPR 250
                                                    His-box II
                               L L                                        F    I
BpΔ6des                    GFSRLWSRFQAWTFVPITSGVFVMAFWLYVLHPSKVFKKKNWEEAFWMLSSHVVRASLIQ 360
OtΔ6des                    GFSKYWLRLQAWTFIPVTSGLV-LLFWMFFLHPSKALKGGKYEELVWMLAAHVIRTWTIK 315
OlΔ6des                    GFSKLWLRLQAWTFVPVTSGMV-LFFWMFVLHPRNALRRKSFEEAAWMFSAHVIRTAVIK 339
MpΔ6des_CCMP1545           GFSRAWARLQAWTFVPVTSGLLVQAFWIYVLHPRQVLRKKNYEEASWMLVSHVVRTAVIK 323
AncΔ6                      GFSRLWLRLQAWTFVPVTSGLLVQLFWMVFLHPRKVLRKKNYEEAAWMLSSHVVRTAVIK 323
MpΔ6des_RCC299             GFSKTWARAQAWTFVPITSGVLVQMFWIYVLHPRQVLRKKNYEEASWMFLSHVVRTAVIK 299
MsΔ6des                    GFSRAWSRAQAWTFVPVTSGLLVQMFWIYVLHPRQVARKKNYEEASWMILSHVLRTATIK 310
                                                                    TMH III
                                        G                                    R
BpΔ6des                    FVHPSNVSFAHAYGLYALSQWIAGMYLFAHFSTSHTHTKVVEFKDHPSWVRYAVEHTVDI 420
OtΔ6des                    AVT--GFTAMQSYGLFLATSWVSGCYLFAHFSTSHTHLDVVPADEHLSWVRYAVDHTIDI 373
OlΔ6des                    AVT--GYSWIASYGLFAATMWASGCYLFAHFSTSHTHLDVVPSDKHLSWVRYAVDHTIDI 397
MpΔ6des_CCMP1545           LAT--GYSWPVAYWWFTFGNWIAYMYLFAHFSTSHTHLPVVPSDKHLSWVNYAVDHTVDI 381
AncΔ6                      AAT--GYSWPQAYGLFALSNWIAGMYLFAHFSTSHTHLDVVPSDKHLSWVRYAVDHTVDI 381
MpΔ6des_RCC299             LAL--GCGTAEAYGWFWVGNWIAYMYLFAHFSTSHTHLDVVPSDKHISWVNYAVDHTVDI 357
MsΔ6des                    YAG--GYSWPVAYLWFSFGNWIAYMYLFAHFSTSHTHLEVVPSDKHISWVNYAVDHTVDI 368
                                  TMH IV
                              N
BpΔ6des                    SPDRAYVNWLMGYLNCQVIHHLFPDMPQFRQPEVSKKFELFARKWGLEYTVMTYGEAWKA 480
OtΔ6des                    DPSQGWVNWLMGYLNCQVIHHLFPSMPQFRQPEVSRRFVAFAKKWNLNYKVMTYAGAWKA 433
OlΔ6des                    NPNNSVVNWLMGYLNCQVIHHLFPDMPQFRQPEVSRRFVPFAKKWNLNYKVLTYYGAWKA 457
MpΔ6des_CCMP1545           DPSRGYVNWLMGYLNCQVIHHLFPDMPQFRQPEVSRRFVPFAKKWGLNYKVLSYYGAWKA 441
AncΔ6                      DPSRGYVNWLMGYLNCQVIHHLFPDMPQFRQPEVSRRFVAFAKKWGLNYKVMTYYGAWKA 441
MpΔ6des_RCC299             NPRNSIVNWLMGYLNCQVIHHLFPDMPQFRQPEVSRRFVAFAEKWNLNYKVLTYYGAWKA 417
MsΔ6des                    DPSKGYVNWLMGYLNCQVIHHLFPDMPQFRQPEVSRRFVAFAKKWNLNYKVLTYYGAWKA 428
                                           His-box III
                                          H
BpΔ6des                    TFKNLNDVGKHYYEEGRVKDKERSSEENAKTKKIK  515
OtΔ6des                    TLGNLDNVGKHYYVHGQHSGKTA------------  456
OlΔ6des                    TFGNLNDVGKHYYVHGSQRVKSKSA----------  482
MpΔ6des_CCMP1545           TFSNLDKVGQHYYVNGKAEKAH-------------  463
AncΔ6                      TFSNLNDVGKHYYVHGKAK----------------  460
MpΔ6des_RCC299             TFSNLDRVGQHYYVNGKAKAH--------------  438
MsΔ6des                    TFTNLDTVGQHYYKHGKAHAH--------------  449
```

**Supplementary Figure 5. The alignment of characterized algal Δ6-desaturases.** The consensus positions are shown in red with the consensus residues identified above the alignment. Positions selected in the consensus of Δ4/5/6-desaturases (Supplementary Fig. 4) are highlighted in grey. Functional motifs are highlighted in yellow and identified below the alignment. Putative MpΔ6des TMHs are underlined and identified below the alignment. AncΔ6, shown here, was not included in the consensus mutagenesis set for algal Δ6-desaturases.

```
MpD6des          MC--PPKTDGRSSPRSPLTRSKSSAEALDAKDASTAPVDLKTLEPHELAA     48
OtD6des          MCVETENNDGIPTVEIAFDGERERAEA-------NVKLSAEKMEPAALAK     43

                                         T    A
MpD6des          TFETRWVRVEDVEYDVTNFKHPGGSVIFYMLANTGADATEAFKEFHMRSL     98
OtD6des          TFARRYVVIEGVEYDVTDFKHPGGTVIFYALSNTGADATEAFKEFHHRSR     93
                                     Heme-binding motif

MpD6des          KAWKMLRALPSRPAEIKRSESEDAPMLEDFARWRAELERDGFFKPSITHV    148
OtD6des          KARKALAALPSRPA--KTAKVDDAEMLQDFAKWRKELERDGFFKPSPAHV    141

                                                                 S
MpD6des          AYRLLELLATFALGTALMYAGYPIIASVVYGAFFGARCGWVQHEGGHNSL    198
OtD6des          AYRFAELAAMYALGTYLMYARYVVSSVLVYACFFGARCGWVQHEGGHSSL    191
                  TMH I                  TMH II              His-box I

MpD6des          TGSVYVDKRLQAMTCGFGLSTSGEMWNQMHNKHHATPQKVRHDMDLDTTP    248
OtD6des          TGNIWWDKRIQAFTAGFGLAGSGDMWNSMHNKHHATPQKVRHDMDLDTTP    241
                                              His-box II

MpD6des          AVAFFNTAVEDNRPRGFSRAWARLQAWTFVPVTSGLLVQAFWIYVLHPRQ    298
OtD6des          AVAFFNTAVEDNRPRGFSKYWLRLQAWTFIPVTSG-LVLLFWMFFLHPSK    290

MpD6des          VLRKKNYEEASWMLVSHVVRTAVIKLATGYSWPVAYWWFTFGNWIAYMYL    348
OtD6des          ALKGGKYEELVWMLAAHVIRTWTIKAVTGFTAMQSYGLFLATSWVSGCYL    340
                       TMH III                    TMH IV

MpD6des          FAHFSTSHTHLPVVPSDKHLSWVNYAVDHTVDIDPSRGYVNWLMGYLNCQ    398
OtD6des          FAHFSTSHTHLDVVPADEHLSWVRYAVDHTIDIDPSQGWVNWLMGYLNCQ    390

                         S
MpD6des          VIHHLFPDMPQFRQPEVSRRFVPFAKKWGLNYKVLSYYGAWKATFSNLDK    448
OtD6des          VIHHLFPSMPQFRQPEVSRRFVAFAKKWNLNYKVMTYAGAWKATLGNLDN    440
                His-box III

MpD6des          VGQHYYVNGKAEKAH-----    463
OtD6des          VGKHYYVHGQ----HSGKTA    456
```

**Supplementary Figure 6. Alignment of MpΔ6des and OtΔ6des.** Mutations from OtΔ6des that were introduced into MpΔ6des are shown in red with the introduced residues identified above the alignment. Positions selected in the consensus of Δ4/5/6-desaturases (Supplementary Fig. 4) are highlighted in grey. Functional motifs are highlighted in yellow and identified below the alignment. Putative MpΔ6des TMHs are underlined and identified below the alignment.

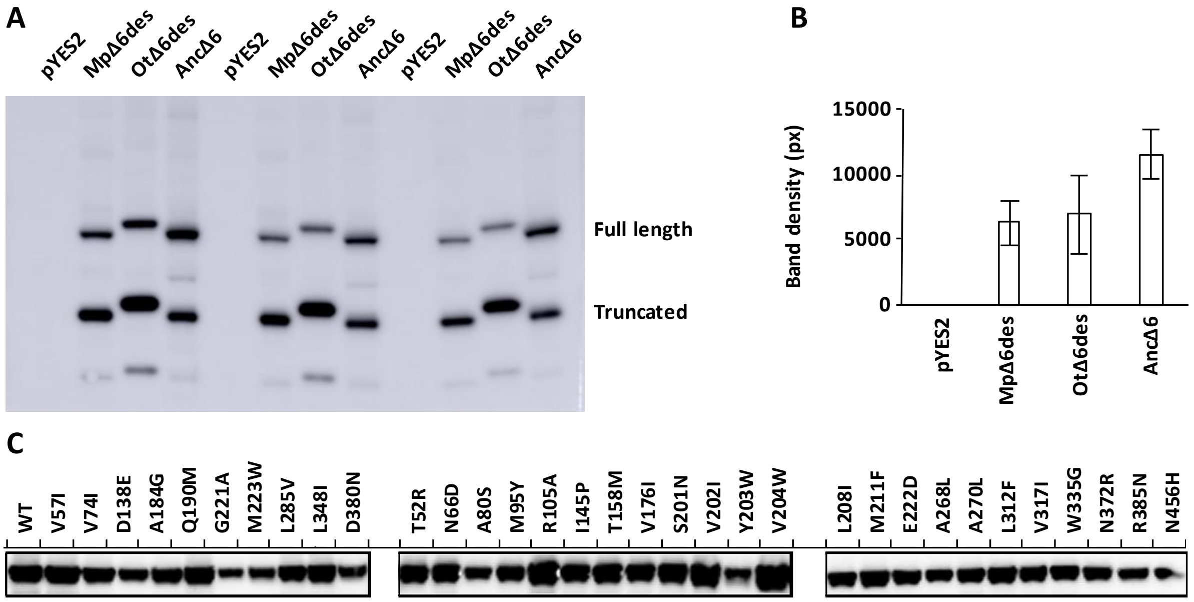


**Supplementary Figure 7. Representative Western blots for Δ6-desaturase expression in *Saccharomyces cerevisiae* S288C.** The expression of Δ6-desaturases was verified by tagging with a hemagglutinin (HA) tag followed by Western blot analysis using anti-HA antibodies. **A.** A triplicate verification of MpΔ6des, OtΔ6des, and AncΔ6 expression with empty pYES2 plasmid as a negative control confirms expression of the desaturases. For all three, expression of a truncated construct still bearing the HA tag is observed. OtΔ6des also demonstrates an increased apparent molecular weight compared to MpΔ6des and AncΔ6 despite its similar theoretical size. This may be due to variations in packing and hydrophobicity. **B.** Relative expression levels of Δ6-desaturases are correlated to band densities. Error shown is the standard deviation of the 3 replicates from panel a). All three desaturases express at significantly higher levels than background observed in the pYES2 negative control. **C.** Western blots of MpΔ6des point mutants from the Δ4/5/6 and Δ6 algal desaturase consensus mutagenesis ensembles show that all tested mutants are readily expressed in *Saccharomyces cerevisiae* S288C.

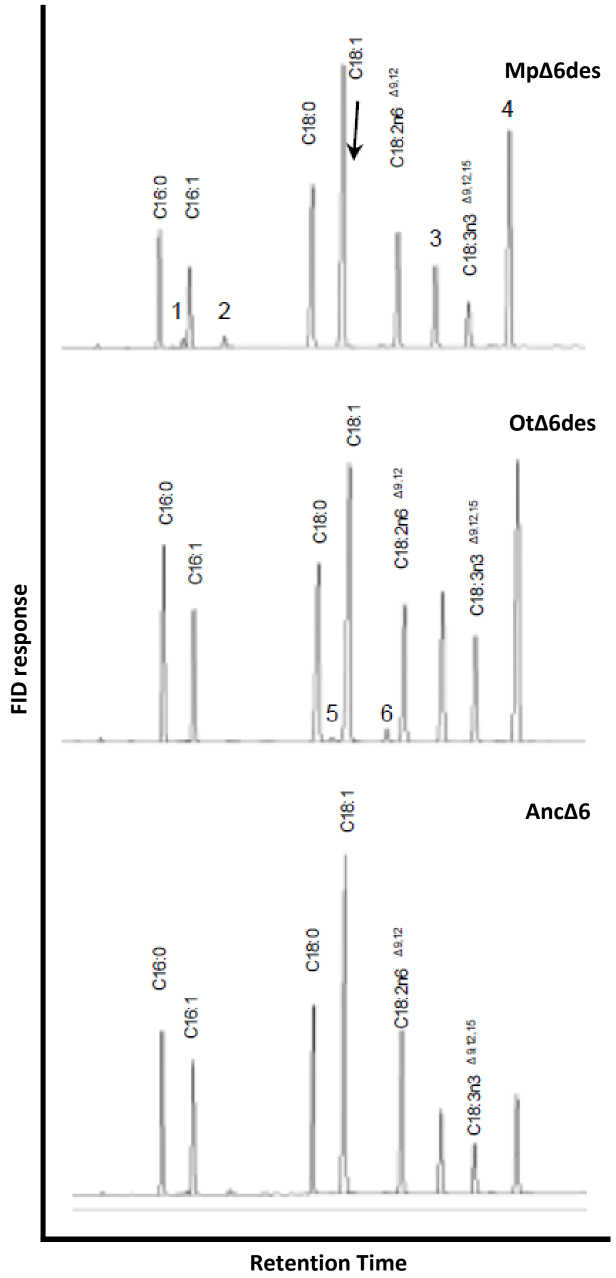


**Supplementary Figure 8. Representative gas chromatography traces for Δ6-desaturase substrate specificity experiments.** Gas chromatography was used to determine Δ6-desaturase substrate specificities using the relative amounts of ω3 and ω6 substrates and product in the total lipid fraction, as indicated by integrated peak volumes for their respective peaks. Peaks labelled by numbers correspond to: 1, C16:1n10$^{\Delta6}$; 2, C16:2n7$^{\Delta6,9}$; 3, C18:3n6$^{\Delta6,9,12}$; 4, C18:4n3$^{\Delta6,9,12,15}$; 5, C18:1n12$^{\Delta6}$; 6, C18:2n9$^{\Delta6,9}$.

### Supplementary Tables

#### Supplementary Table 1. Sequences included in the desaturase phylogenetic tree.

| Organism | GenBank Accession | Substrate | Regiospecificity[a] |
|---|---|---|---|
| *Isochrysis galbana* | AAV33631.1 | ω3-DPA 28%<br>ω6-DPA 13% | Δ4 |
| *Isochrysis galbana* | AFF27583.1 | ω3-DPA 34% (highest)<br>ω6-DPA NO DATA | Δ4 |
| *Isochrysis* sp. GB-2012 | AFJ74710.1 | DPA 79.8%,<br>Active with both ω3 and ω6. | Δ4 |
| *Isochrysis galbana* CCMM5001 | AFB82637 | No Data | Δ6 |
| *Bathycoccus prasinos* | CCO20439 | ND | Δ6 |
| *Micromonas pusilla* CCMP1545 | XP_003056992 | LA 4.9%<br>ALA 63.0% | Δ6/8 |
| *Mantoniella squamata* | CAQ30479 | ALA 34%<br>LA 0.3% | Δ6 |
| *Micromonas* sp. RCC299 | XP_002502445 | NO DATA | Δ6 |
| *Ostreococcus tauri* | XP_003082578 | LA 73%<br>ALA 71% | Δ6 |
| *Ostreococcus lucimarinus* CCE9901 | DAA34893 | LA 6.6%<br>ALA 38.8% | Δ6 |
| *Ostreococcus lucimarinus* CCE9901 | XP_001421073 | NO DATA | Δ6 |
| *Leishmania braziliensis* | XP_001562610 | No DATA | Δ5 |
| *Leishmania mexicana* | XP_003872262 | No DATA | Δ5 |
| Codon optimized *Leishmania major* | AEI59766 | DGLA ω6 5%<br>ETA ω3 5% | Δ5 |
| *Leishmania* major strain Friedlin | XP_001681021 | No DATA | Δ5 |
| *Leishmania donovani* | XP_003858558 | No DATA | Δ5 |
| *Leishmania infantum* JPCM5 | XP_001463326 | No DATA | Δ5 |
| *Perkinsus marinus* ATCC 50983 | XP_002765309 | No DATA | Δ5/6 |
| *Perkinsus marinus* ATCC 50983 | XP_002765314 | No DATA | Δ5/6 |
| *Perkinsus marinus* | ABF58685 | C20 DGLA and EPA both active.<br>Slight ω6 preference | Δ5 |
| *Thraustochytrium aureum* | BAK08911 | DGLA ω6 22.9%<br>ETA ω3 19.9% | Δ5 |
| *Thraustochytrium sp* | AAM09687.1 | ω6 22:4(7,10,13,16)<br>ω3 22:5(7,10,13,16,19) | Δ4 |
| *Thraustochytrium* sp. FJN-10 | ACD03117 | DGLA ω6 56.4%<br>No ω3 data | Δ5 |
| *Isochrysis galbana* | AEA72469.1 | DGLA ω6 active | Δ5 |

| | | | |
|---|---|---|---|
| *Isochrysis galbana* | AFD22890.1 | No DATA | Δ5 |
| *Rebecca salina* | ABL96295.1 | DGLA ω6 96.7%<br>ETA ω3 99.2% | Δ5 |
| *Aureococcus anophagefferens* | EGB03053 | No DATA | Δ6-like |
| *Aureococcus anophagefferens* | EGB08313 | No DATA | Δ6-like |
| *Salpingoeca rosetta* | EGD73438 | No DATA | Δ5 |
| *Monosiga brevicollis* MX1 | XP_001743517 | No DATA | Δ5 |
| *Capsaspora owczarzaki* ATCC 30864 | EFW47040 | No DATA | Δ8 |
| *Amphimedon queenslandica* | XP_003385370 | No DATA | FAD2-like |
| *Clonorchis sinensis* | GAA50852 | No DATA | FAD2-like |
| *Octopus vulgaris* | AEK20864 | No DATA | Δ5 |
| *Crassostrea gigas* | EKC30965 | No DATA | FAD2 |
| *Haliotis discus hannai* | ADK38580 | No DATA | Δ5 |
| *Haliotis discus hannai* | ADK12703 | No DATA | Δ5 |
| *Crassostrea gigas* | EKC33620 | No DATA | FAD2 |
| *Pseudomonas brassicacearum* | AEA70230 | No DATA | HP |
| *Taeniopygia guttata* | XP_002194906 | No DATA | HP |
| *Gallus gallus* | XP_426408 | No DATA | HP |
| *Meleagris gallopavo* | XP_003206387 | No DATA | HP |
| *Anolis carolinensis* | XP_003224187 | No DATA | HP |
| *Taeniopygia guttata* | XP_002194862 | No DATA | HP |
| *Scyliorhinus canicula* | AEY94454 | DGLA ω6 29%<br>ETA ω3 55% | Δ5 |
| *Mus musculus* | BAE32539 | No DATA | HP |
| *Mus musculus* | NP_666206 | C20:2n-9 is active | Δ5 FAD1 |
| *Mus musculus* | AAH26848 | No DATA | HP |
| *Mus musculus* | AAH22139 | No DATA | HP |
| *Rattus norvegicus* | AAG35068 | DGLA is active.<br>ω3 activity no data | Δ5 |
| *Rattus norvegicus* | NP_445897 | Activity is shown | Δ5 |
| *Rattus norvegicus* | AEX15917 | No DATA | FAD1 |
| *Canis lupus familiaris* | XP_540914 | No DATA | HP |
| *Ailuropoda melanoleuca* | EFB20951 | No DATA | HP |
| *Otolemur garnettii* | XP_003798788 | No DATA | HP |
| *Bos taurus* | XP_612398 | No DATA | HP |
| *Nomascus leucogenys* | XP_003274097 | No DATA | HP |
| *Nomascus leucogenys* | XP_003274097 | No DATA | HP |
| *Homo sapiens* | AAF29378 | DGLA is active | Δ5 |

| | | | |
|---|---|---|---|
| *Homo sapiens* | BAC11182 | No DATA | HP |
| *Homo sapiens* | AAF70457 | DGLA and ETA is active | Δ5 |
| *Homo sapiens* | BAB55103 | No DATA | HP |
| *Homo sapiens* | BAC11229 | No DATA | HP |
| *Homo sapiens* | AAH07846 | No DATA | HP |
| Synthetic human gene | AAX29339 | No DATA | FAD1 |
| *Homo sapiens* | NP_037534 | No DATA | FAD1 |
| *Pan troglodytes* | JAA30224 | No DATA | FAD1 |
| *Homo sapiens* | BAD96626 | No DATA | FAD1 |
| *Homo sapiens* | BAB55173 | No DATA | HP |
| *Pan paniscus* | XP_003828511 | No DATA | HP |
| *Anolis carolinensis* | XP_003224167 | No DATA | HP |
| *Gallus gallus* | XP_421052 | No DATA | HP |
| *Taeniopygia guttata* | XP_002194926 | No DATA | HP |
| *Saccoglossus kowalevskii* | XP_002739666 | No DATA | HP |
| *Mus musculus* | BAE32659 | No DATA | HP |
| *Cricetulus griseus* | XP_003513228 | No DATA | HP |
| *Mus musculus* | BAA95044 | No DATA | HP |
| *Mus musculus* | BAC37908 | No DATA | HP |
| *Mus musculus* | NP_068690 | No DATA | FAD3 |
| *Mus musculus* | BAC26393 | No DATA | HP |
| *Pangasianodon hypophthalmus* | AFN21428 | No DATA | HP |
| *Danio rerio* | Q9DEX7 | ALA, LA, SDA and GLA. Δ6 activity higher than Δ5. ω3 preference. | Δ5/6 |
| *Danio rerio* | NP_571720 | No DATA | Desaturase |
| *Salmo salar* | NP_001117014 | No DATA | Δ5 |
| *Gallus gallus* | NP_001153900 | No DATA | HP |
| *Scyliorhinus canicula* | AEY94455 | LA 57%.<br>ALA 73% | Δ6 |
| *Xenopus laevis* | NP_001083680 | No DATA | HP |
| *Xenopus (Silurana) tropicalis* | NP_001120262 | No DATA | HP |
| *Xenopus laevis* | NP_001086853 | No DATA | HP |
| *Ceratodon purpureus* | CAB94993 | LA | Δ6 |
| *Cunninghamella echinulata* | ABA06503 | LA | Δ6 |
| *Conidiobolus obscurus* | AEA07665 | LA 16%.<br>ALA 15% | Δ6 |
| *Mortierella alpina* | ADE06661 | No DATA | Δ5 |
| *Mortierella alpina* | AAF08685 | LA | Δ6 |

| | | | |
|---|---|---|---|
| *Mortierella alpina* | CAE53093 | No DATA | Δ6 |
| *Mortierella alpina* | ABN69091 | No DATA | Δ6 |
| *Mortierella alpina* | BAA85588 | LA and ALA | Δ6 |
| *Mortierella alpina* | AAF08685 | LA | Δ6 |
| *Umbelopsis isabellina* | AAG38104 | No DATA | Δ6 |
| *Umbelopsis isabellina* | AAL73948 | No DATA | Δ6 |

[a] HP: Hypothetical Protein

### Supplementary Table 2 – Relative activity and specificity of MpΔ6des variants. [a]

| Group | Protein | Normalized ω3 activity [b] | Normalized ω6 activity [c] | Relative ω3/ω6 specificity [d] |
|---|---|---|---|---|
| | WT | 1.00 ± 0.07 | 1.00 ± 0.14 | 1.00 ± 0.16 |
| Algal Δ4/5/6-desaturases | V57I | 1.01 ± 0.02 | 1.07 ± 0.06 | 0.95 ± 0.06 |
| | V74I | 0.87 ± 0.04 | 0.44 ± 0.04 | 1.98 ± 0.11 |
| | D138E | 1.01 ± 0.01 | 1.02 ± 0.02 | 0.99 ± 0.02 |
| | A184G | 0.90 ± 0.07 | 0.57 ± 0.10 | 1.57 ± 0.19 |
| | Q190M | 0.88 ± 0.01 | 0.29 ± 0.01 | 3.03 ± 0.03 |
| | G221A | 0.89 ± 0.03 | 0.57 ± 0.06 | 1.57 ± 0.10 |
| | M223W | 0.49 ± 0.04 | 0.18 ± 0.02 | 2.73 ± 0.11 |
| | L285V | 1.01 ± 0.02 | 1.06 ± 0.04 | 0.96 ± 0.04 |
| | L348I | 0.33 ± 0.05 | 0.04 ± 0.01 | 8.33 ± 0.43 |
| | D380N | 0.76 ± 0.01 | 0.22 ± 0.01 | 3.53 ± 0.05 |
| Algal Δ6-desaturases | T52R | 0.88 ± 0.01 | 0.64 ± 0.03 | 1.37 ± 0.04 |
| | N66D | 0.95 ± 0.06 | 0.85 ± 0.15 | 1.12 ± 0.18 |
| | A80S | 1.01 ± 0.06 | 0.93 ± 0.13 | 1.08 ± 0.16 |
| | M95Y | 0.84 ± 0.01 | 0.53 ± 0.01 | 1.60 ± 0.03 |
| | R105A | 1.04 ± 0.05 | 1.10 ± 0.13 | 0.94 ± 0.13 |
| | I145P | 1.09 ± 0.05 | 1.22 ± 0.15 | 0.89 ± 0.14 |
| | T158M | 1.04 ± 0.01 | 1.12 ± 0.03 | 0.93 ± 0.03 |
| | V176I | 1.07 ± 0.04 | 1.30 ± 0.13 | 0.82 ± 0.11 |
| | S201N | 1.08 ± 0.03 | 1.22 ± 0.34 | 0.89 ± 0.30 |
| | V202I | 1.06 ± 0.04 | 1.25 ± 0.13 | 0.85 ± 0.12 |
| | Y203W | 1.07 ± 0.03 | 1.30 ± 0.11 | 0.82 ± 0.10 |
| | V204W | 0.99 ± 0.10 | 0.93 ± 0.20 | 1.07 ± 0.24 |
| | L208I | 1.05 ± 0.08 | 1.11 ± 0.17 | 0.95 ± 0.18 |
| | M211F | 1.06 ± 0.05 | 1.35 ± 0.31 | 0.79 ± 0.25 |
| | E222D | 1.06 ± 0.09 | 1.56 ± 0.27 | 0.68 ± 0.20 |
| | A268L | 1.08 ± 0.08 | 1.07 ± 0.16 | 1.01 ± 0.18 |
| | A270L | 1.20 ± 0.01 | 1.63 ± 0.30 | 0.74 ± 0.22 |
| | L312F | 1.17 ± 0.08 | 1.14 ± 0.14 | 1.03 ± 0.17 |
| | V317I | 1.19 ± 0.07 | 1.21 ± 0.17 | 0.98 ± 0.18 |
| | W335G | 0.78 ± 0.05 | 0.39 ± 0.07 | 2.02 ± 0.17 |
| | N372R | 0.87 ± 0.01 | 0.58 ± 0.05 | 1.49 ± 0.07 |
| | R385N | 0.84 ± 0.01 | 0.55 ± 0.01 | 1.51 ± 0.02 |
| | N456H | 0.85 ± 0.02 | 0.75 ± 0.16 | 1.14 ± 0.18 |
| OtΔ6des | S73T | 0.99 ± 0.01 | 0.97 ± 0.04 | 1.03 ± 0.05 |
| | M78F | 1.02 ± 0.01 | 1.16 ± 0.03 | 0.88 ± 0.03 |
| | N196S | 0.98 ± 0.01 | 0.92 ± 0.03 | 1.06 ± 0.04 |
| | D406S | 0.87 ± 0.01 | 0.40 ± 0.01 | 2.15 ± 0.02 |
| | S-N/A-L | 1.05 ± 0.04 | 1.57 ± 0.14 | 0.67 ± 0.10 |
| | E-D/A-L | 1.04 ± 0.04 | 1.72 ± 0.13 | 0.60 ± 0.08 |
| | S-N/E-D | 0.97 ± 0.03 | 1.17 ± 0.09 | 0.83 ± 0.08 |
| | V3 | 1.08 ± 0.02 | 2.04 ± 0.13 | 0.53 ± 0.07 |

[a] All values given as the mean of 3 replicates ± 1 S.D.
[b] The ω3 substrate (ALA) conversion of the mutant normalized by the ALA conversion of the wild type
[c] The ω6 substrate (LA) conversion of the mutant normalized by the LA conversion of the wild type
[d] The ALA/LA conversion ratio of the mutant normalized by the ALA/LA conversion ratio of the wild type

**Supplementary Table 3. Summary of experimentally characterized algal Δ6-desaturase specificities.**

| Enzyme (GenBank ID) | Organism | Conversion efficiency (%) | | Geographic origin | Referenc e |
|---|---|---|---|---|---|
| | | LA (ω6) | ALA (ω3) | | |
| MpΔ6des (XP_003056992) | *Micromonas pusilla* CCMP1545 | 4.9 | 63.0 | English Channel (Atlantic ocean) | [4] |
| MsΔ6des (CAQ30479) | *Mantoniella squamata* | 0.3 | 34.0 | Atlantic Ocean | [29] |
| OtΔ6des (XP_003082578) | *Ostreococcus tauri* | 73.0 | 71.0 | Mediterranean Sea | [32] |
| OlΔ6des (DAA34893) | *Ostreococcus lucimarinus* CCE9901 | 6.6 | 38.8 | Pacific Ocean | [31] |
